# The Potential of the Primitive: a Network Analysis of Early Arthropod Evolution

**DOI:** 10.1101/2023.01.03.522586

**Authors:** Agustín Ostachuk

## Abstract

2

It is often thought that the primitive is simpler, and that the complex is generated from the simple by some process of self-assembly or self-organization, which ultimately consists of the spontaneous and fortuitous collision of elementary units. This idea is included in the Darwinian theory of evolution, to which is added the competitive mechanism of natural selection. To test this view, we studied the early evolution of arthropods. Twelve groups of arthropods belonging to the Burgess Shale, Orsten Lagerstätte, and extant primitive groups were selected, their external morphology abstracted and codified in the language of network theory. The analysis of these networks through different network measures (network parameters, topological descriptors, complexity measures) was used to carry out a PCA and a hierarchical clustering, which allowed us to obtain an evolutionary tree with distinctive/novel features. The analysis of centrality measures revealed that these measures decreased throughout the evolutionary process, and led to the creation of the concept of evolutionary developmental potential. This potential, which measures the capacity of a morphological unit to generate changes in its surroundings, is concomitantly reduced throughout the evolutionary process, and shows that the primitive is not simple but has a potential that unfolds during this process. This means for us the first evolutionary empirical evidence of our theory of evolution as a process of unfolding.

**Teaser:** Primitive is not simple but has a potential that unfolds throughout the evolutionary process.

## 4 Introduction

It is usually thought that the primitive is simpler. This line of thought also ensures that the complex is generated from the simple by some process of self-assembly or self-organization, with diverse variations and conceptual tonalities. This concept developed contemporaneously in the field of cybernetics, although its origins can be traced back to the concept that a natural end must not only be organized, but must also be self-organized, expounded in Kant’s *Kritik der Urteilskraft* of 1790 [1], and more rudimentarily in the Greek materialism developed first by Leucippus and Democritus, and later by Epicurus and Lucretius, who believed that everything is generated and can be generated by the spontaneous and fortuitous collision of elementary particles or atoms [2].

The so-called principle of “self-organization” was proposed, as such, by William Ross Ashby in 1947 [3], and then underwent successive reformulations and subsequent developments. Thus, for example, another cybernetician, Heinz von Foerster, formulated the principle of “order from noise” in 1960 [4], the biophysicist Henri Atlan then based on this concept to develop his principle of “complexity from noise” in 1972 [5], and the chemist Ilya Prigogine (together with Isabelle Stengers) later formulated a similar principle called “order out of chaos” in 1984 [6]. The concept of self-organization assumes that order is produced spontaneously through a process of local interactions between parts from an initially disordered system. This process is facilitated by random disturbances (“noise”) that allow the system to explore different states until arriving and being attracted by a stable state called attractor.

Similarly, in the field of biology it is assumed that the evolution of organisms occurred through a random and aleatory process of organic modifications (in Darwin’s time morphological changes, currently genetic mutations), whose temporal accumulation enabled the generation of new species. What was added in this case, compared to the previous case, was a mechanism proposed by Darwin called “natural selection” [7]. According to this mechanism, only the species best adapted to their environments could survive (“the survival of the fittest”, according to Herbert Spencer), in a context where there was constant competition between species and “struggle for existence”, with a clear influence and origin in liberal economy [8].

This line of thought has greatly influenced the theories that try to explain the structure and form of the most primitive arthropods and crustaceans. There is a general historical consensus that the primitive arthropod or crustacean, *Urarthropod* or *Urcrustacean*, consisted of a simple organism. Thus, for example, Hessler & Newman (1975) [9], considering Cephalocarida, Leptostraca (Malacostraca) and Notostraca (Branchiopoda) as the most primitive representatives of the crustaceans, speculated that the most likely ancestor of the crustaceans, based on morphological similarity, were the trilobites. Trilobites present a very simple general organization characterized by serial homology and the presence of stenopodous limbs, that is, a serial, repetitive and linear morpho-topological structure. Something similar happened when the class Remipedia was discovered [10]. Their long bodies with many similar segments without specialization and their serially homonomous nonphyllopodous trunk limbs were quickly interpreted as evidence of their primitiveness, and as the likely extant crustacean with the most primitive body plan [11]. Remipedia was considered the sister group of Cephalocarida, as we saw, another crustacean class generally regarded as one of the most primitive [12]. Nowadays, the situation has changed and they are not considered a primitive crustacean but a sister group of insects [13]. However, the general situation has not changed and what is understood by primitive arthropod or crustacean, and the structural and morphological organization attributed to it, remains largely the same.

In this manner, both the principle of self-organization, used by cybernetics, and the principle of natural selection, used by biology, assume that the evident increase in the complexity of organisms occurred throughout evolutionary history, has occurred by a random and spontaneous process of interactions between simpler elements. In short, these theories assume and maintain that the complex can arise from the simpler. We have developed a whole theory of evolution based on the opposite principle: the complex cannot arise from the simpler [14]. We have already given the fundamental reason for founding this principle: the simpler does not have the necessary information for the generation of something more complex than itself. From a logical point of view, the proposition that the evolution from a unicellular organism to a primate occurred by a mere process of random and spontaneous interactions between elementary parts does not seem to have solid support. In a previous work we had found evidence for our theory in the ontogenetic and metamorphic development of the crab [15]. In this work we will present evidence of the theory in the early evolution of arthropods. We developed network models of some of the most important organisms for the study of the origin and early evolution of arthropods: fossils from the Burgess Shale and Orsten Lagerstätte, and extant crustaceans generally considered to be primitive and original in the subphylum. Several network measures (network parameters, topological descriptors and complexity measures) were evaluated, and with them a PCA and Hierarchical clustering were carried out. This allowed the construction of a hypothetical evolutionary tree. This tree posed a scenario in which the evolution of arthropods is marked by an early bifurcation into two evolutionary branches: a left or large branch (networks greater than 400 nodes), and a right or small branch (networks less than 400 nodes). Predictably, *Yohoia* and *Canadaspis* were placed as primitive arthropods, as the originators of the left branch. Surprisingly, *Triops* (Branchiopoda: Notostraca) was placed as the successor of this branch, before other organisms usually considered more primitive, such as *Rehbachiella* and *Marrella*; and *Branchinecta* (Branchiopoda: Anostraca) was located as the originator of the right branch, before other organisms usually considered more primitive, such as *Waptia* and *Olenoides* (Trilobita). The evolutionary tree allowed at the same time to establish a rationale and to understand the behavior of the evaluated network measures. These measures had two basic patterns: ascending bifurcation and descending bifurcation. A large part of the network measures had an ascending bifurcation pattern, that is, they increased throughout the evolutionary process, which were characterized as extensive complexity measures in the previous work [15]. On the other hand, a group of network measures had a descending bifurcation pattern, decreasing their values throughout the evolutionary process. To study in greater detail the meaning of these results, the organisms were studied at the level of their morpho-topological structure by measuring different centrality measures. The results showed that most of these measures had a descending bifurcation pattern. Several of these centrality measures measure the degree of influence and power of an actor within the network. This allowed us to create the concept of *evolutionary developmental potential*, which measures the ability or capacity of a morphological structure to generate changes in its area of influence. Visualization of arthropod networks based on these measures showed that primitive organisms possessed a high concentration of evolutionary developmental potential throughout the body, while later organisms only had a high concentration in the head. Consequently, we concluded that the evolutionary developmental potential decreases throughout the evolutionary process, determining a lower capacity to generate changes in later organisms. We interpret these empirical results as clear evidence for the theory of evolution as a process of unfolding that we have recently published [14].

## 5 Materials and Methods

### 5.1 The Arthropod Network Model

Arthropod networks were built according to the same principles as crustacean networks [15, 16], and can be considered an extension and broadening of those principles to the phylum Arthropoda. The distinctive characteristic of these groups is that they are formed by segments that are articulated in different ways among themselves. This morphological pattern can be translated into network theory as nodes connected to each other by edges. In this manner, the different morphological units of arthropods, clearly identifiable, distinguishable and delimited, generically named as segments or articles, were considered nodes. The physical connections between these elements, articulated or non-articulated, were considered edges.

In order to study the early evolution of arthropods, the most representative groups of arthropods considered primitive due to their morphological characteristics were chosen, that is, fossils from the Burgess Shale and the Orsten Lagerstätte. To study the relationship between these extinct primitive arthropods and arthropods existing today, some of the groups of living crustaceans considered to be the most primitive and original of the subphylum were included in the analysis. Table 1 details the list of arthropod groups included in the analysis, and the references used for the detailed and thorough study of their external morphology.

**Table 1:**
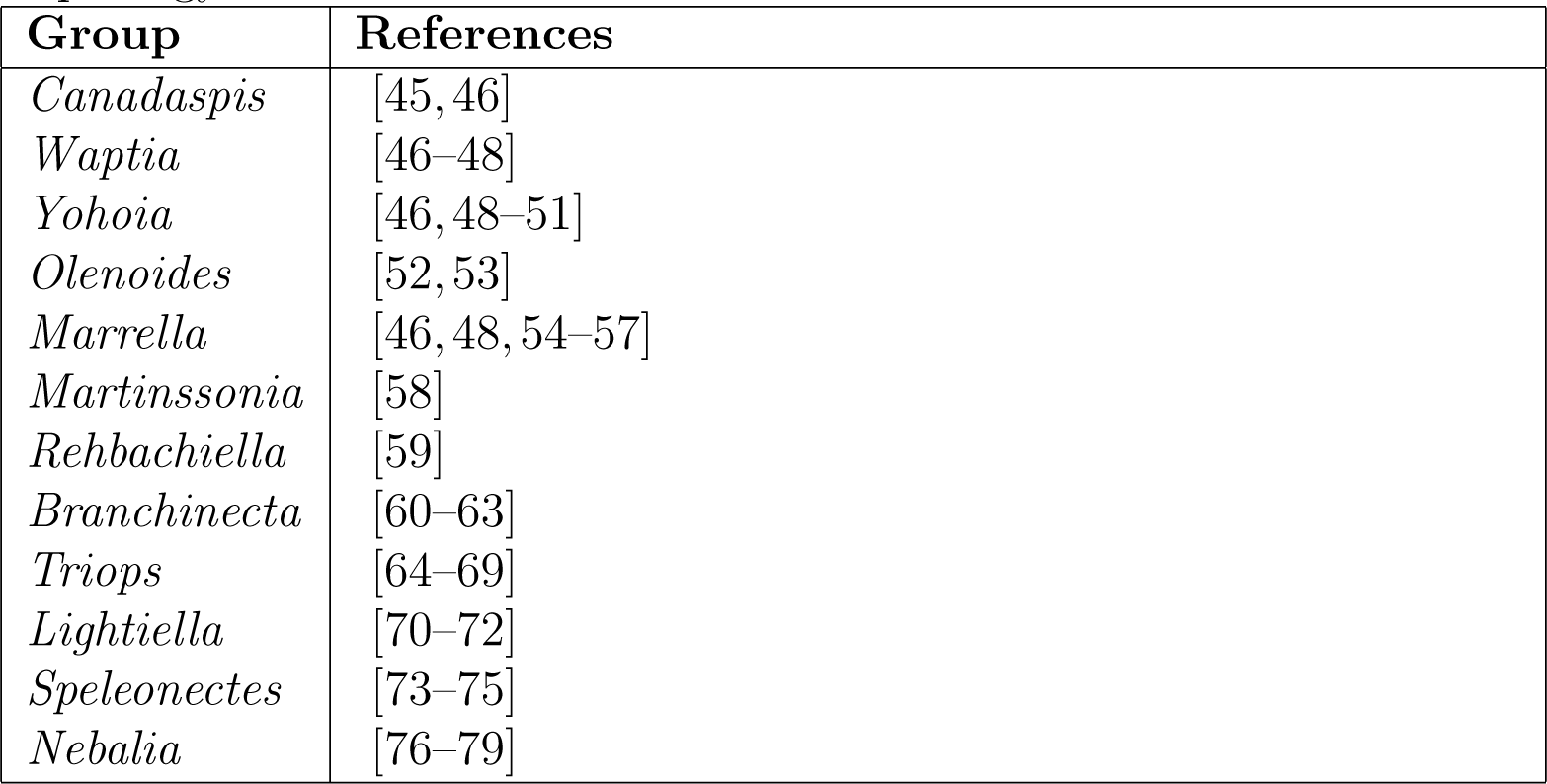
Groups of extinct and extant arthropods selected for the present work and the corresponding references used for the study of their external morphology

The detailed and systematic study of the morphology of arthropods, their different morphological units and connections between them, allows the creation of a blueprint or design plan of the network. This diagram is then translated into the computer language of network theory by preparing the adjacency matrix, which is a two-axis table made up of all the morphological units of the animal, and where the connection between two units is represented by the number 1 and the lack of connection by the number 0. In this manner, the adjacency matrix has a dimension of *N* × *N* , where *N* represents the total number of nodes in the network. This matrix, typically an object of *csv* file extension, is then used to build the actual network, typically an object of *graphml* file extension. The generation of the network from the adjacency matrix was carried out using the R programming language [17, 18], and the *igraph* network analysis package [19]. Networks were displayed and spatialized using the ForceAtlas 2 layout algorithm [20], belonging to the network visualization software *Gephi* [21].

### 5.2 Network parameter analysis

Once the corresponding network object has been obtained, it is possible to proceed to the analysis of its elemental characteristics. For a first approximation to the characterization of the network properties, the following parameters were calculated: number of nodes, number of edges, diameter, radius, average path length, average degree, average clustering coefficient, density. All these parameters can be obtained with the *igraph* package.

### 5.3 Topological descriptors

To carry out a deeper and more detailed analysis of arthropod networks, a battery of topological descriptors initially developed for the identification and discrimination of molecules and chemical structures was evaluated. Recently, these descriptors have been used for studies of morphological evolution, showing enormous analytical power to reveal topological properties of biological networks [15, 16].

These descriptors measure different topological properties of a network. Some descriptors consist of measures based on distance. Thus, for example, the Wiener index, the first topological descriptor developed and perhaps the best known, consists of the sum of the distances between each pair of nodes in the network [22]. Another well-known distance-based descriptor is the Balaban J index. This index is obtained using the distance matrix, and calculating the distance sum for each node of the network, which is why they are also called distance degrees [23]. Other descriptors are based on entropy measures, such as the Bonchev index [24] and the Bertz complexity index [25]. Another set of descriptors consists of measures based on the eigenvalues. Within this group are the Estrada index, the Laplacian Estrada index, the Energy index and the Laplacian Energy index [26]. Finally, other descriptors are based on other invariants, such as the Zagreb index [27], the Randű connectivity index [28], the Complexity index B and Normalized Edge complexity [29]. The latter is also called connectedness, and represents the quotient between the sum of all node degrees and the number of edges of the complete graph.

### 5.4 Complexity measures

Some of the topological descriptors mentioned above can be considered as complexity measures. However, many of them do not meet an important requirement to be able to compare the complexity between different networks: they are not normalized complexity measures. Consequently, comparison between networks of different sizes is difficult. For this case, there is another package of complexity measures that are normalized. This group of measures assumes that the most complex networks have an intermediate number of edges, since this allows the existence in them, unlike what happens in complete networks, of characteristic intricate internal structures, such as modular, hierarchical structures and specific functional regions.

Of the three existing groups of complexity measures, in this work we used two: Product measures and Entropy measures. Within the first group are Medium Articulation (*MAg*), Efficiency complexity (*Ce*) and Graph index complexity (*Cr*). *MAg* is basically the product between redundancy and mutual information, followed by normalization (Kim 2008). *Ce* measures how efficiently a network exchanges information using the inverse shortest path lengths [30]. *Cr* is based on the properties of the largest eigenvalue of the adjacency matrix, the index *r* [31]. The three measures were taken into account in this work. For its part, from the second group, we evaluated the measure called Offdiagonal complexity (*OdC*). *OdC* measures diversity in the node-node link correlation matrix [32].

### 5.5 Principal Component Analysis (PCA)

Once all the networks parameters, topological descritptors and complexity measures were obtained, all of them were used to carry out a Principal Component Analysis (PCA), with the idea of revealing hidden patterns derived from the structure and topology of the networks, and their relationship with the different topological and complexity measures. The PCA was carried out using the *prcomp* function of the R programming language. The visualization of the individuals and variables for the three main dimensions of the PCA was carried out using the *factoextra* package [33].

### 5.6 Hierarchical clustering

Hierarchical cluster analysis (HCA) is a clustering method for grouping objects based on their similarity. In agglomerative clustering, each observation is initially considered as a cluster (leaf), and then the most similar clusters are successively merged until a single cluster (root) is obtained. The result of a hierarchical clustering is a tree-based representation of the objects, i.e. a dendrogram. First, the data containing the results obtained for the different topological and complexity measures were scaled, using the *scale* function of the R programming language. Second, the dissimilarity matrix was calculated to measure the degree of (dis)similarity between the networks. For this, the *dist* function of the R programming language was used to compute the Euclidean distance between the networks. Finally, a linkage function was applied to group, from the distance information, pairs of objects into clusters based on their similarity. This is an iterative process that is repeated until all the objects are linked in a hierarchical tree. This was done with the *hclust* function of the R programming language, choosing the Ward method (*ward.D*2), which minimizes the total within-cluster variance: in each step the clusters that are merged are those with the smallest between-cluster distance. The result of the HCA was visualized using the *factoextra* package, which allows visualizing the clustering with different graphical representations of the dendrogram (horizontal/vertical, circular, phylogenic). In order to compare results, another clustering method was tested, Hierarchical K-Means Clustering, which combines the best of k-means clustering and hierarchical clustering. For this, the *hkmeans* function from the *factoextra* package was used.

### 5.7 Centrality measures

Centrality measures were calculated using the *igraph* [19] and *netrankr* [34] packages. For its part, the visualization of the networks with the different centrality measures was carried out using the *tidygraph* [35] and *ggraph* [36] packages. To do this, it is necessary to first transform the network, usually from *igraph* class, to *tbl* class.

## 6 Results

### 6.1 Early arthropod evolution: group selection and network visualization

For the study of early arthropod evolution, we chose 12 groups of animals that we considered the most representative in order to best cover the wide spectrum and diversity of groups since the appearance of the phylum Arthropoda (specifically since the appearance of the upper stem-group euarthropods, following the phylogenetic tree developed by Legg *et al.* (2013) [37]), including Trilobitomorpha as a representative of Artiopoda, until the appearance of the members of Pancrustacea usually considered the most primitive (that is, we sought to cover the part of the phylogenetic tree developed by Legg *et al.* (2013) [37] included in Fig. 4B, 4C and 4H). We considered that this number of representatives allowed us to cover the wide spectrum and diversity of forms that were generated in early arthropod evolution, and to carry out a meticulous and detailed study of this early evolution. In other words, it allowed us the development of a general, qualitative and theoretical research, and a specific, quantitative and empirical research at the same time. We also sought to include the most characteristic and iconic groups of two of the most important sites of primitive arthropods: Burgess Shale and Orsten Lagerstätte. Thus, the groups included in our study were the following: *Canadaspis*, *Waptia*, *Yohoia*, *Olenoides* (Trilobita), *Marrella*, *Martinssonia*, *Rehbachiella*, *Branchinecta* (Branchiopoda: Anostraca), *Triops* (Branchiopoda: Notostraca), *Lightiella* (Cephalocarida), *Speleonectes* (Remipedia) and *Nebalia* (Malacostraca: Leptostraca) (Table 1).

Fig. 1 shows the networks resulting from the analysis of the morphological structure of these arthropods and their abstraction in the form of nodes connected by edges. In general terms, the networks ranged from a size of 250 (*Martinssonia*) to 929 nodes (*Triops*). The characteristics and properties of these networks will be discussed later.

**Figure 1.**
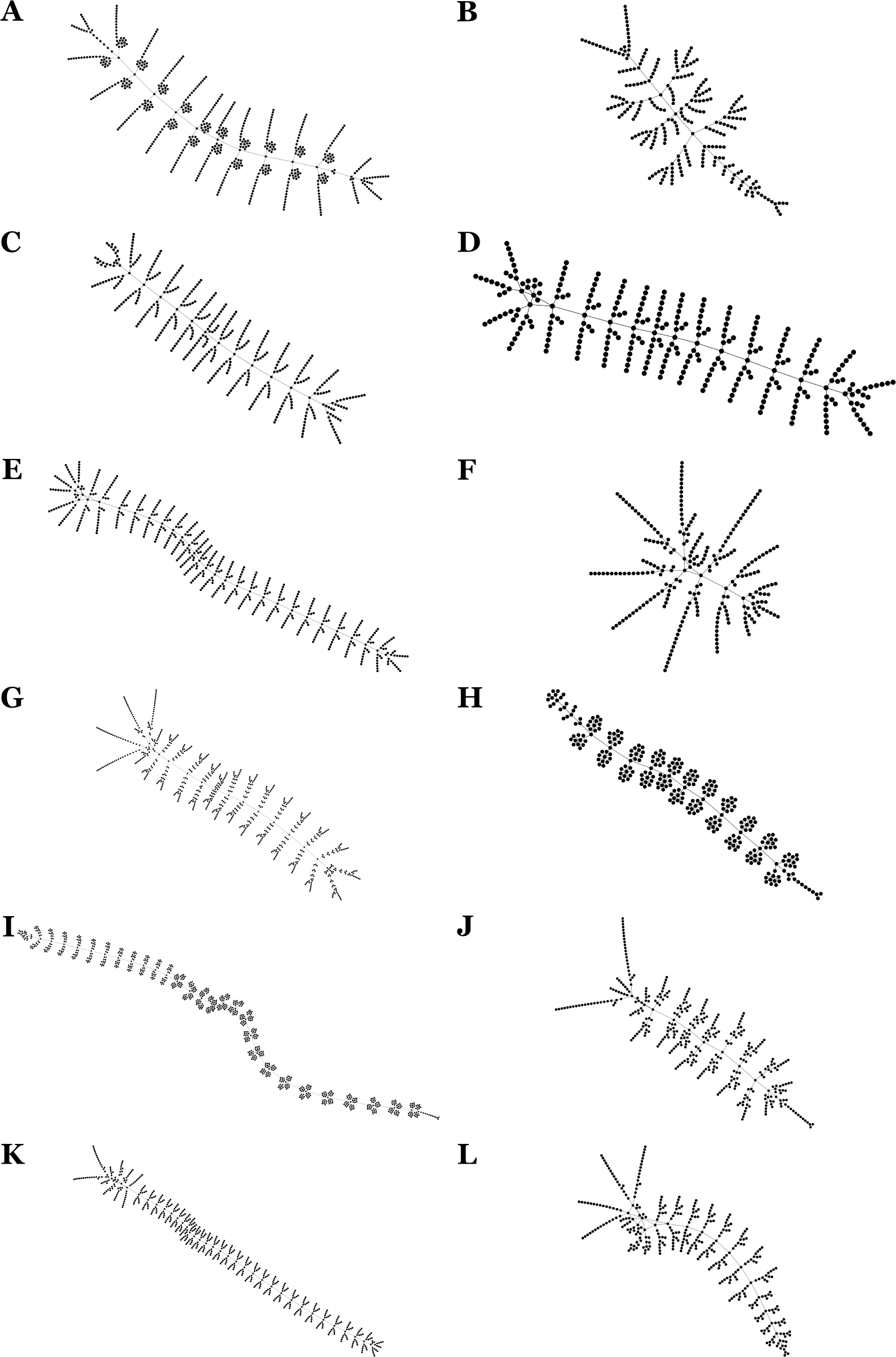
Arthropod networks. The networks resulting from the abstraction of the external morphology of the selected arthropods and their coding to network theory are shown using the *Gephi* network visualization software. Networks were displayed and spatialized using the ForceAtlas 2 layout algorithm. Node size is the same in all cases, and the difference in the size of the nodes that you see in the figure is due to a difference in scale. (A) *Canadaspis*, (B) *Waptia*, (C) *Yohoia*, (D) *Olenoides*, (E) *Marrella*, (F) *Martinssonia*, (G) *Rehbachiella*, (H) *Branchinecta*, (I) *Triops*, (J) *Lightiella*, (K) *Speleonectes*, (L) *Nebalia*.

### 6.2 Principal Component Analysis (PCA)

The results obtained for the network parameters, topological descriptors and complexity measures did not give a recognizable pattern ordered according to the phylogenetic tree used as a reference in this work [37]. As a consequence, we decided to carry out a PCA to try to reveal hidden and not entirely evident patterns, using the three groups of network measures just mentioned. This analysis revealed surprising and interesting evolutionary processes, which will become clearer with the deep analyzes carried out later (hierarchical clustering and centrality measures).

It is worth mentioning that the result of the PCA seems to be very robust, since it was carried out only with the topological descriptors and the results obtained were very similar, almost identical, to those obtained in the PCA analyzed below, in which the topological descriptors, network parameters and complexity measures were included.

#### 6.2.1 Network analysis

From a topological point of view, the PCA seemed to distribute the networks into 5 different topological regions (Fig. 2 and 3). A first region (Region 1) was characterized by being located in the positive zone of dimension 2 and 3, occupying approximately the central position with respect to dimension 1. The networks corresponding to *Canadaspis*, *Yohoia* and *Waptia* were located in this region, the latter being located in the positive zone of dimension 1. A second region (Region 2) was characterized by being located in the negative zone of dimension 1 and 2, occupying approximately the central region with respect to dimension 3. In this region the network corresponding to *Triops* was found. A third region (Region 3) was characterized by being located in the positive zone of dimension 1 and the negative zone of dimension 2, also occupying approximately the central region with respect to dimension 3. The network corresponding to *Branchinecta* was found in this region. Region 3 was then located in the opposite zone with respect to dimension 1 than Region 2. In turn, Region 2 and 3 were located in the opposite zone with respect to dimension 2 than Region 1. For its part, a fourth region (Region 4) was characterized by being located in the negative zone of dimension 1 and 3, and the positive zone of dimension 2. The networks corresponding to *Rehbachiella*, *Marrella* and *Speleonectes* were found in this region. Finally, a fifth region (Region 5) was characterized by being located in the positive zone of dimension 1 and 2, and the negative zone of dimension 3. The networks corresponding to *Olenoides*, *Martinssonia*, *Nebalia* and *Lightiella* were found in this region. Region 5 was then located on the opposite zone with respect to dimension 1 than Region 4, such that Region 5 shared the positive zone of dimension 1 with Region 3, and Region 4 shared the negative zone of dimension 1 with Region 2. For their part, Region 4 and 5 occupied the opposite region with respect to dimension 3 than Region 1, while they coincided with respect to dimension 2.

**Figure 2.**
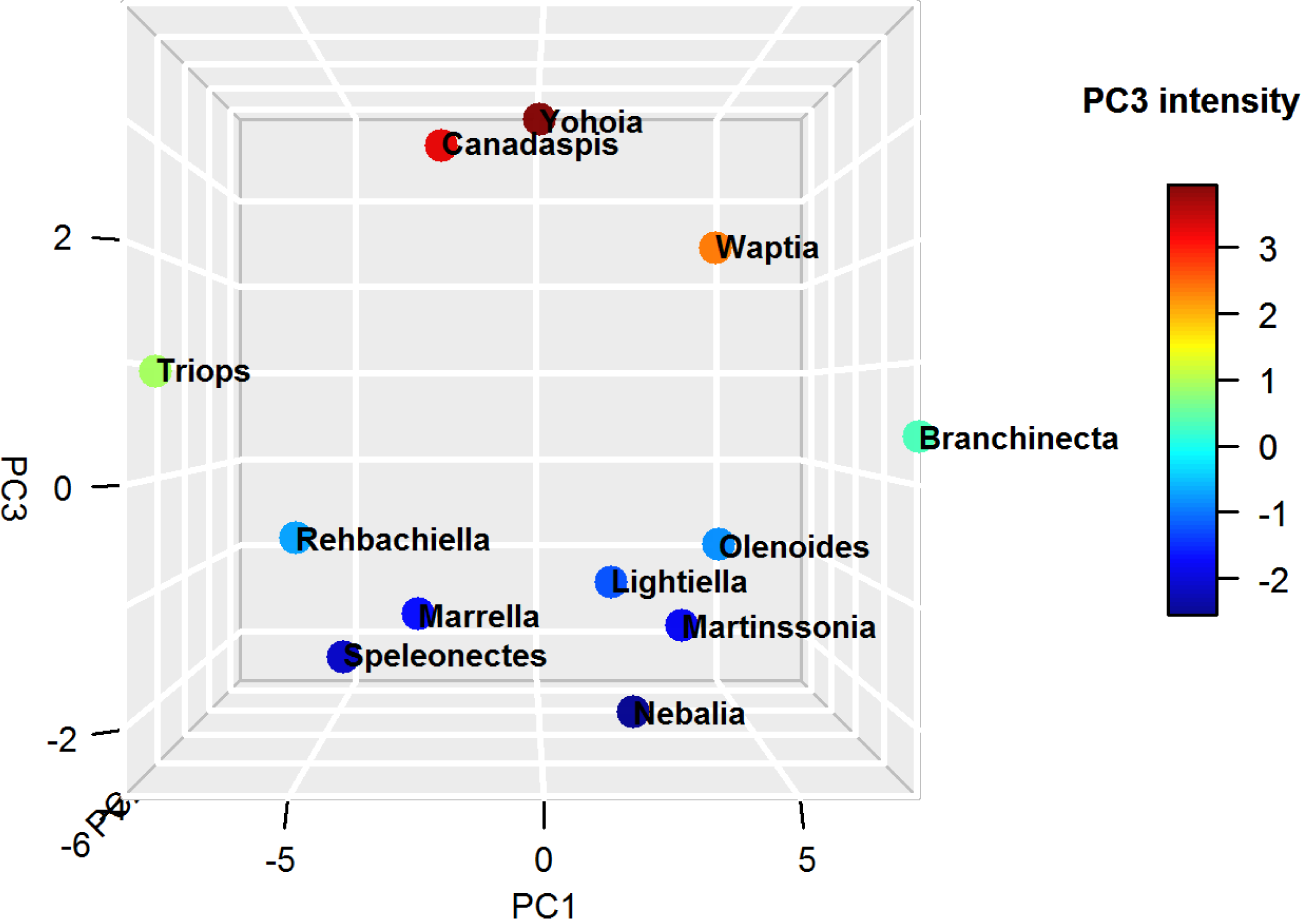
Principal Component Analysis (PCA). Distribution of arthropod networks in a three-dimensional space formed by the first three principal components (orientation: PC3 vs. PC1). Dot color varies according to PC3 intensity (see lateral scale).

**Figure 3.**
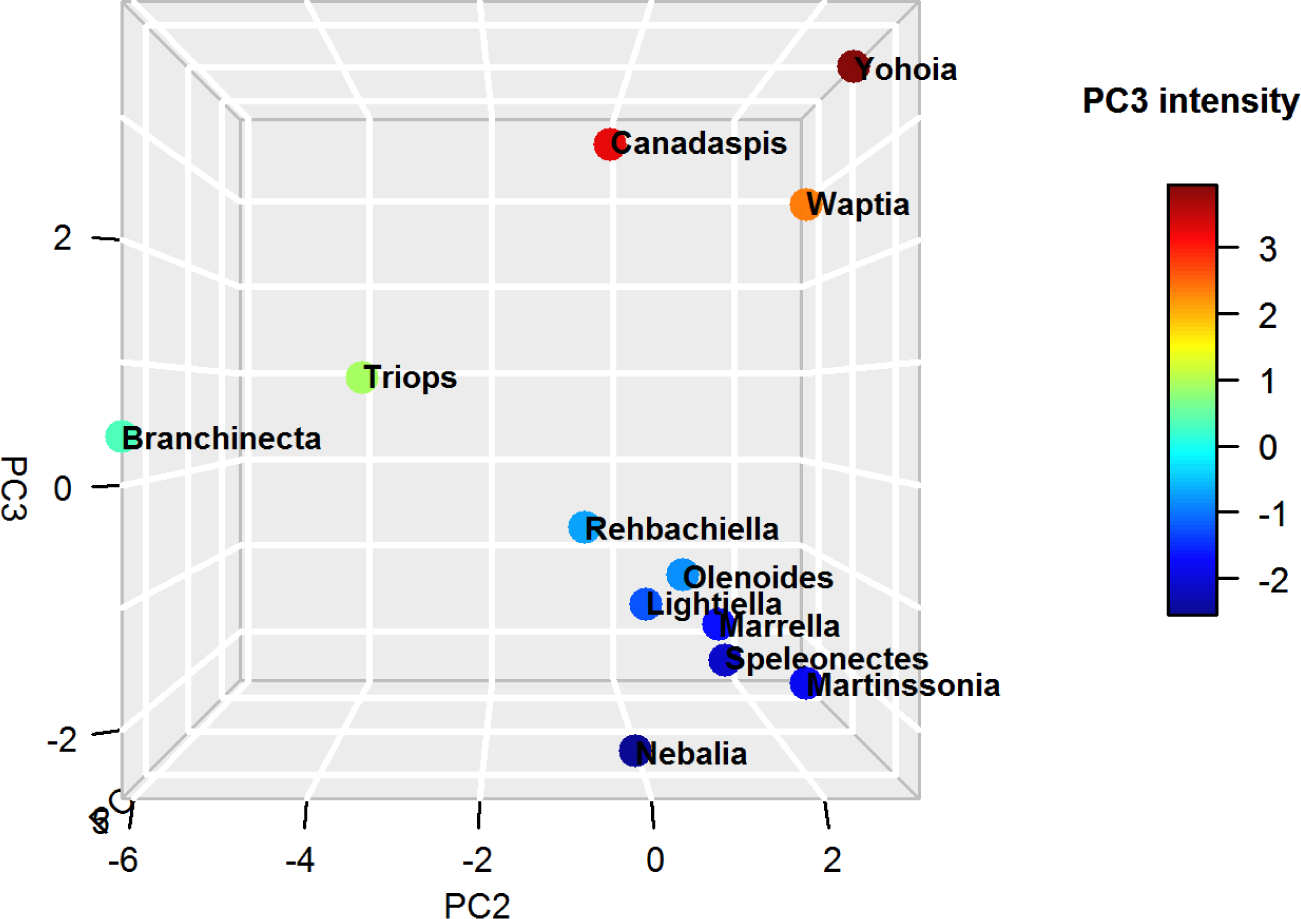
Principal Component Analysis (PCA). Distribution of arthropod networks in a three-dimensional space formed by the first three principal components (orientation: PC3 vs. PC2). Dot color varies according to PC3 intensity (see lateral scale).

The richest information seemed to be found in dimensions 1 and 3 of the PCA (Fig. 2). Dimension 1 explained 60.6 % of the variation, while dimension 3 explained 12.2 % of the variation. For their part, the first three dimensions as a whole explained 90.6 % of the variation. Fig. 2 seems to show a scenario where a group of original arthropods, formed by *Canadaspis*, *Yohoia*, and perhaps also *Waptia*, gave rise to two large evolutionary branches: (1) left or large branch (networks greater than 400 nodes), led by *Triops*, and made up of the largest arthropods (*Rehbachiella*, *Marrella*, *Speleonectes*); and (2) right or small branch (networks less than 400 nodes), led by *Branchinecta*, and formed by the smallest arthropods (*Olenoides*, *Martinssonia*, *Nebalia*, *Lightiella*). According to the results of this analysis, dimension 3 would be determining the temporality of the evolutionary process, while dimension 1 would be indicating the spatiality/diversification of this process.

If this evolutionary scenario is correct, two quite surprising events stand out and are revealed as novel with respect to the currently existing bibliography: (1) the existence of a very early bifurcation of arthropods, and (2) a prominent and decisive role of *Triops* and *Branchinecta* in this early process of bifurcation. These results and evidences, if verified, would provide a novel and, up to now, unexplored view of early arthropod evolution. The orders Anostraca and Notostraca, belonging to class Branchiopoda, are considered primitive crustaceans, but clearly not at the level of basal arthropods. This privileged position of *Triops*, as a representative of Notostraca, and *Branchinecta*, as a representative of Anostraca, in the early evolution of arthropods is reaffirmed and underlined by dimension 2 of the PCA (Fig. 3). Basically, this dimension separates these two groups from the rest of the other groups of arthropods. Later on, we will try to explain the structural cause of this difference, as well as the structural cause that characterizes the most primitive arthropods according to the present analysis (*Canadaspis*, *Yohoia*, *Waptia*). In other words, the structural changes that would be taking place along dimensions 2 and 3 of this analysis will be investigated. In the next section we will analyze the contribution of the different network measures to the first three dimensions of the PCA.

#### 6.2.2 Network measures analysis

An important number of network measures (Group 1) contributed mainly to the negative axis of dimension 1, with different degrees of contribution to the negative axis of dimension 2, while they contributed very little to dimension 3 (region: PC1 negative, PC2 center to negative, PC3 center) (Fig. 4 and 5). A subgroup of network measures were almost aligned with the negative axis of dimension 1 (Randć connectivity index, Average distance, Mean distance deviation, Energy), while another subgroup contributed moderately to the negative axis of dimension 2 (Nodes, Edges, Wiener index, Centralization, Eccentricity, Zagreb index 1, Bertz complexity index, Bonchev index 2, Konstantinova index, Information layer index), and a third subgroup contributed highly to the negative axis of dimension 2 (Estrada index, Harary index, Laplacian energy, Zagreb index 2).

**Figure 4.**
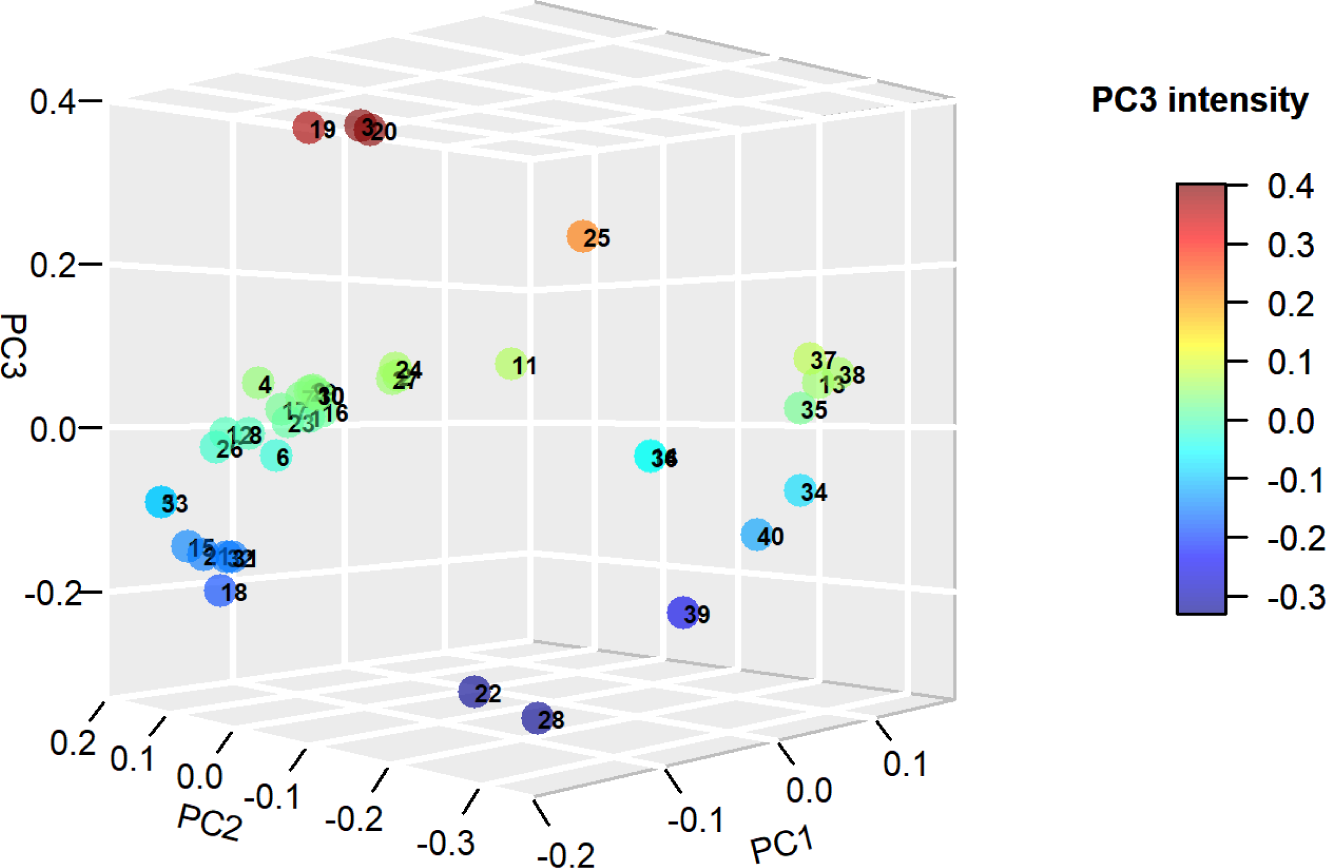
Principal Component Analysis (PCA). Distribution of network measures in a three-dimensional space formed by the first three principal components. Dot color varies according to PC3 intensity (see lateral scale). Network measures: (1) Wiener index, (2) Harary index, (3) Balaban J index, (4) Mean distance deviation, (5) Compactness, (6) Eccentricity, (7) Centralization, (8) Average distance, (9) Konstantinova index, (10) Zagreb index 1, (11) Zagreb index 2, (12) Randű connectivity index, (13) Complexity index B, (14) Normalized edge complexity, (15) Bonchev index 1, (16) Bonchev index 2, (17) Bertz complexity index, (18) Radial centric information index, (19) Balaban-like information index 1, (20) Balaban-like information index 2, (21) Graph vertex complexity index, (22) Edge equality, (23) Information layer index, (24) Estrada index, (25) Laplacian Estrada index, (26) Energy, (27) Laplacian energy, (28) Spectral radius, (29) Nodes, (30) Edges, (31) Diameter, (32) Radius, (33) Average path length, (34) Average degree, (35) Average clustering coefficient, (36) Density, (37) *MAg*, (38) *Ce*, (39) *Cr*, (40) *OdC*.

**Figure 5.**
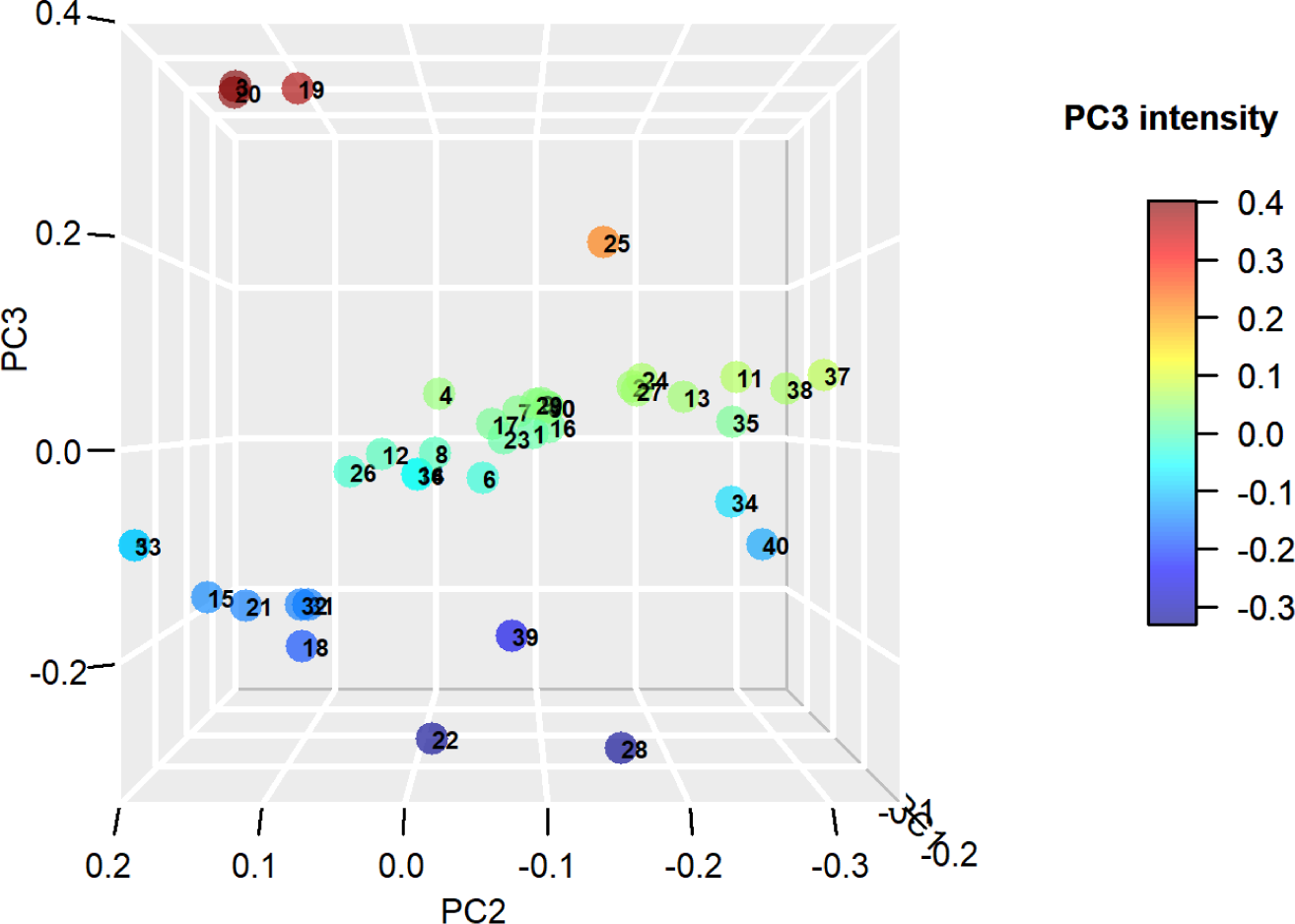
Principal Component Analysis (PCA). Distribution of network measures in a three-dimensional space formed by the first three principal components (orientation: PC3 vs. PC2). Dot color varies according to PC3 intensity (see lateral scale). See Fig. 4 for network measures’ numbering.

The group located in the region opposite to the previous group (Group 2) was characterized by contributing to the positive axis of dimension 1, and contributing very little to the other two dimensions (region: PC1 positive, PC2 center, PC3 center). The network measures Density, Normalized edge complexity and the complexity measure *Cr* were found in this group. The latter, unlike the other two, not only contributed to the positive axis of dimension 1, but also to the negative axis of dimension 3. Group 1 and 2 were the groups that were projected and essentially determined dimension 1 of the PCA, in its negative and positive direction, respectively.

Another group of network measures (Group 3) not only contributed to the negative axis of dimension 1, but at the same time to the positive axis of dimension 2 and to the negative axis of dimension 3 (region: PC1 negative, PC2 positive, PC3 negative). These are the network measures Diameter, Radius, Average path length, Graph vertex complexity, Compactness, Bonchev index 1 and Radial centric information index.

The group that was somehow arranged in the opposite region to the previous group (Group 4) was characterized by contributing to a great extent to the negative axis of dimension 2, with a variable contribution to the positive axis of dimension 1, and little contribution to dimension 3 (region: PC1 center to positive, PC2 negative, PC3 center). The following network measures could be found in this group: Average degree, Average clustering coefficient, Complexity index B, *MAg*, *Ce*, *OdC*. Most of the complexity measures used were found in this group. Group 3 and 4 were the groups that were projected and essentially determined dimension 2 of the PCA, in its positive and negative direction, respectively.

A very important group (Group 5), for reasons that we will see and analyze later, were the network measures characterized by contributing to the positive axis of dimension 2 and 3 (region: PC1 center, PC2 positive, PC3 positive). The network measures Balaban J index, Balaban-line information index 1 and Balaban-like information index 2 were found in this group. The Balaban-like information index 1, unlike the other two, had a median contribution to the negative axis of the dimension 1.

The group that was somehow arranged in the opposite region to the previous group (Group 6) was characterized by contributing essentially to the negative axis of dimension 3, with a very low contribution to dimension 1 and 2 (region: PC1 center, PC2 center, PC3 negative). The network measures Edge equality and Spectral radius were found in this group. The second had a moderate contribution to the negative axis of dimension 2. Groups 5 and 6 were the groups that were projected and essentially determined dimension 3 of the PCA, in its positive and negative direction, respectively.

### 6.3 Hierarchical Clustering (HC)

The PCA provided very valuable information and seemed to reveal an evolutionary pattern determined by dimension 3 as temporality and dimension 1 as spatiality/diversification, in which *Canadaspis*, *Yohoia* and *Waptia* appeared as the most primitive arthropods, surprisingly followed by *Triops* and *Branchinecta*, inaugurating two well differentiated evolutionary lineages. To confirm or refute this interpretation of the PCA, a Hierarchical Clustering (HC) was carried out based on the results of the network measures. This involved first scaling these results, then calculating the degree of (dis)similarity between the networks, and finally grouping the networks into clusters according to their degree of similarity, following an agglomeration method. The HC used in this work is equivalent to the one known as Unweighted Pair-Group Method using Arithmetic average (UPGMA), although using *ward.D*2 as agglomeration method. The difference between the two is that the branch lengths change.

The result of the HC is shown in the dendrograms of Figures 6 and 7, using two different types of dendrograms, “rectangle” and “phylogenic”, respectively. The correlation between the cophenetic distances and the original distance data (cophenetic correlation coefficient) was 0.685. Using another clustering method, Hierarchical K-Means Clustering, which combines the best of k-means clustering and hierarchical clustering, gave exactly the same results. All this is an indication that the results are solid and robust.

**Figure 6.**
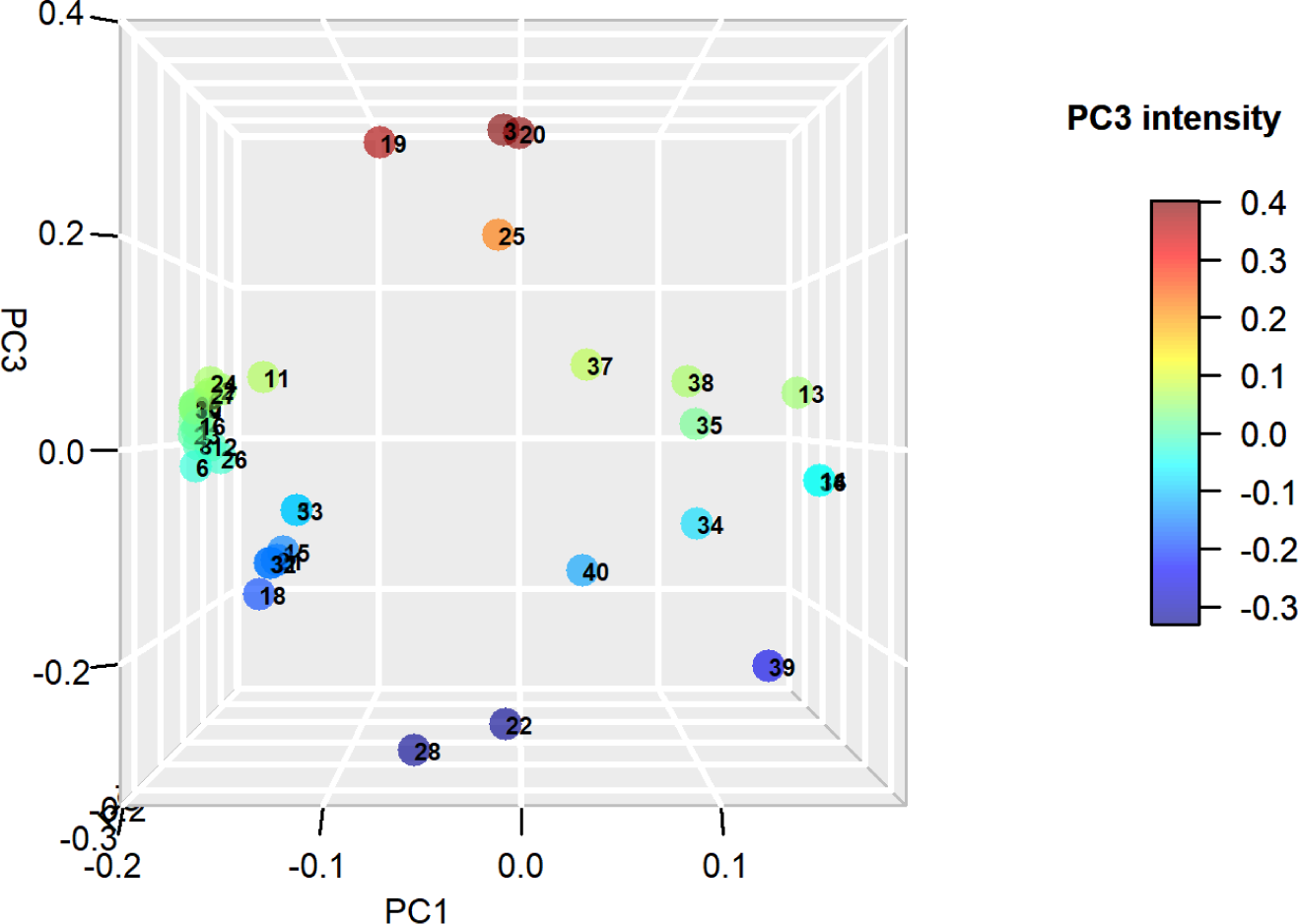
Principal Component Analysis (PCA). Distribution of network measures in a three-dimensional space formed by the first three principal components (orientation: PC3 vs. PC1). Dot color varies according to PC3 intensity (see lateral scale). See Fig. 4 for network measures’ numbering.

**Figure 7.**
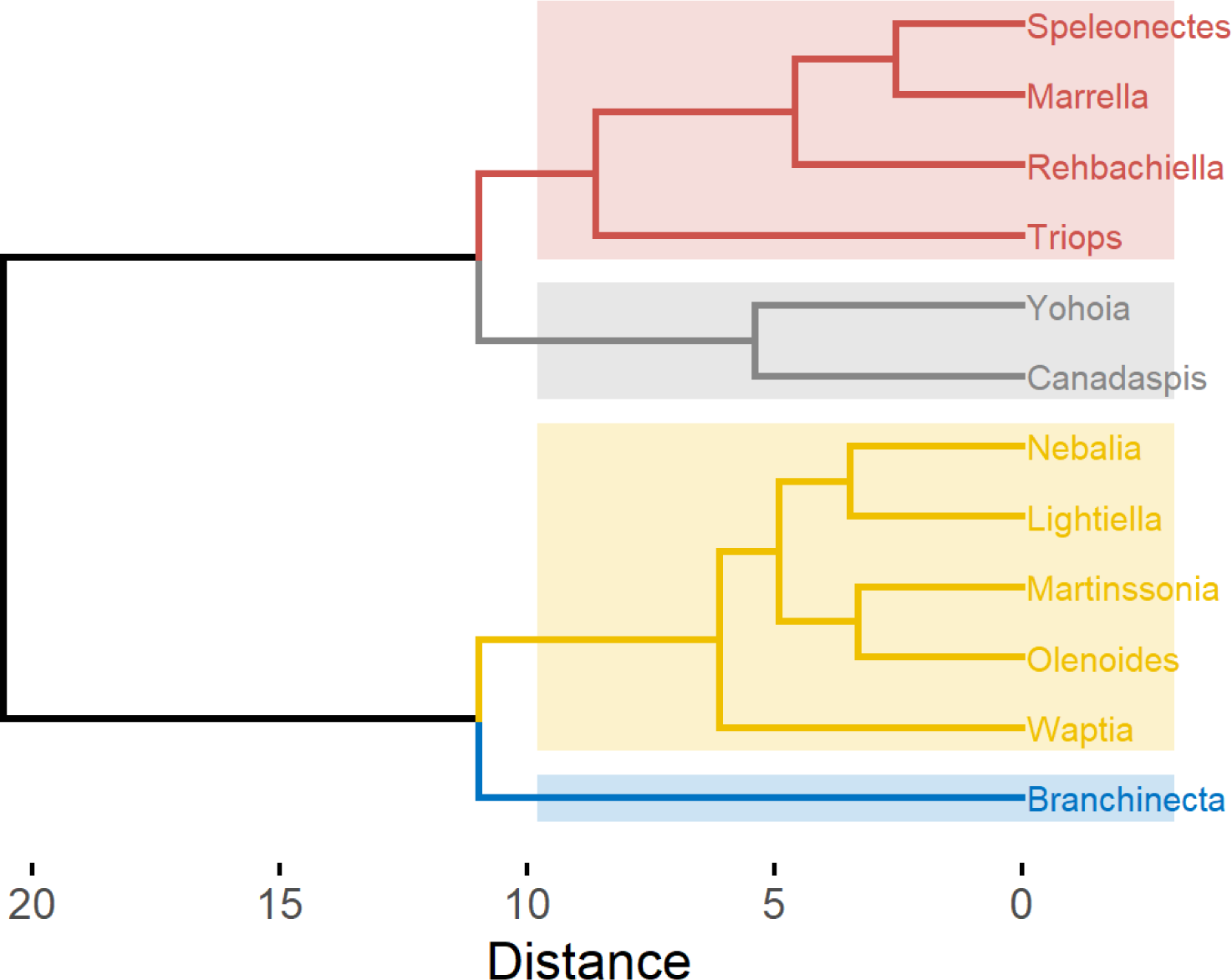
Hierarchical Clustering (HC). Horizontal dendrogram of arthropod networks carried out by agglomerative hierarchical clustering using Ward’s method (*ward.D2*). The dendrogram was divided into 4 clusters, shown in different colors, based on the degree of similarity.

The result of the HC largely confirms the qualitative analysis performed on the PCA, with some interesting differences. Firstly, it places the origin of arthropods at an intermediate point between the *Canadaspis*-*Yohoia* pair, and *Branchinecta*. Secondly, *Branchinecta* is placed as the originator of what we have called the right or small branch, while *Canadaspis* and *Yohoia*, their common ancestor to be more precise, are placed as the originators of the left or large branch. Thirdly, *Waptia* then appears as the first specimen of the branch originating from *Branchinecta*, while *Triops* appears as the first specimen of the branch originating from *Canadaspis*-*Yohoia*. In this manner, the statistical analysis carried out by means of hierarchical clustering locates the origin of arthropods not so much in *Canadaspis* and *Yohoia*, as the PCA result suggested, but in an ancestor of these two and of *Branchinecta*, probably more similar to the latter than the first two. As a consequence of this, *Waptia* is still considered a primitive group, but posterior and derived from *Branchinecta*. The location of *Canadaspis* and *Yohoia* as two of the most primitive arthropods is not surprising, and it is what is usually believed of these Burgess Shale specimens. However, the positions of *Branchinecta* and *Triops* close to the point of origin, and giving rise to two early evolutionary branches of arthropods (in the case of *Triops* following *Canadaspis* and *Yohoia*), is an intriguing and novel result. In subsequent sections we will see what morphological and topological characteristics would explain the primitiveness of these organisms, and what modifications of these characteristics would occur throughout the evolutionary process.

The representation of the hierarchical clustering in the form of a heatmap (Fig. 9) provides a more visual analysis of the previously analyzed results, at the same time as it provides a hierarchical clustering of the network measures. We can see in this format that there is a clear topological difference between the networks of organisms belonging to the right or small branch, and those belonging to the left or large branch. For example, while the networks of the right branch present negative values of the network measures that we had grouped in Group 1 in the PCA, the networks of the left branch present positive values for these measures. Practically the same occurs with the network measures that we had grouped into Group 3 in the PCA. On the other hand, the opposite occurs with the measures grouped in Group 2: while the networks of the right branch present positive values for these network measures, the networks of the left branch present negative values.

**Figure 8.**
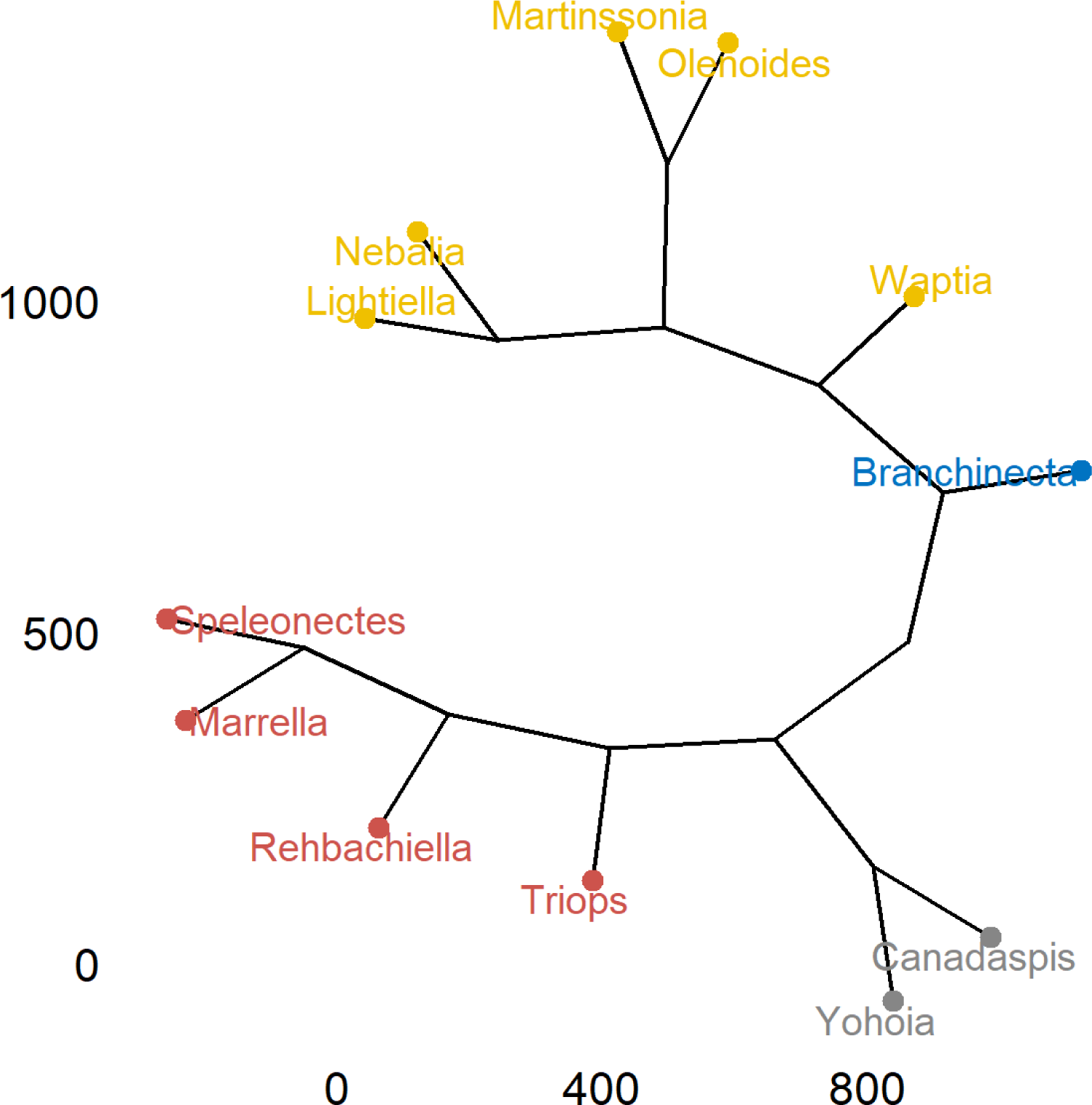
Hierarchical Clustering (HC). Phylogenic dendrogram of arthropod networks carried out by agglomerative hierarchical clustering using Ward’s method (*ward.D2*). The dendrogram was divided into 4 clusters, shown in different colors, based on the degree of similarity.

**Figure 9.**
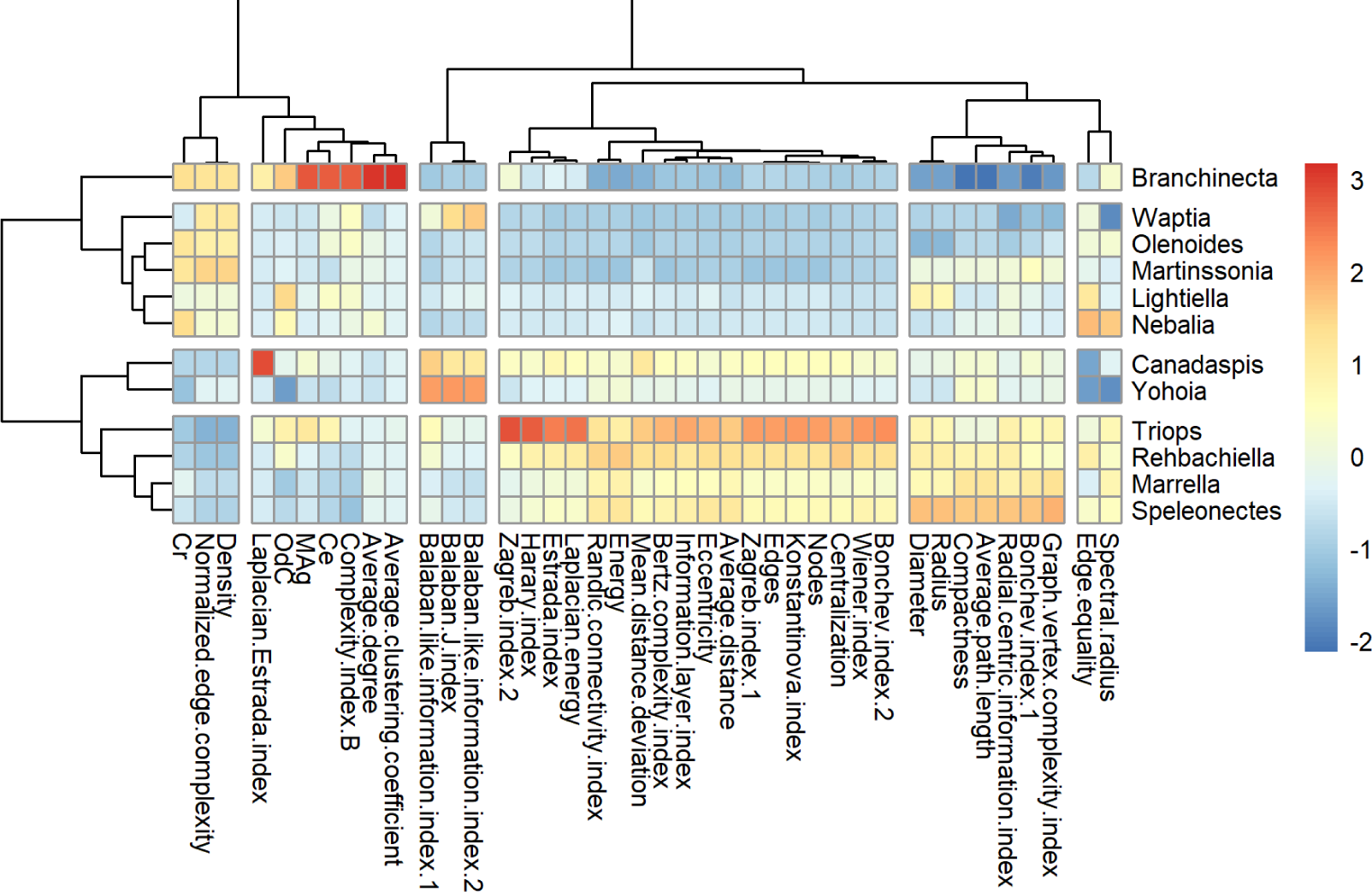
Hierarchical Clustering (HC). Heatmap of arthropod networks and network measures carried out by agglomerative hierarchical clustering using Ward’s method (*ward.D2*). The dendrogram of arthropod networks was divided into 4 clusters, while the dendrogram of network measures was divided into 6 clusters, based on the degree of similarity. Color represents the value of the scaled measures (see lateral scale).

The hierarchical clustering practiced on the network measures coincides exactly with the division of groups that we had carried out qualitatively in the PCA. The same 6 groups defined in the PCA are the same 6 clusters into which the hierarchical clustering can be divided, each of them being formed by the same network measures as those previously defined (with the exception of the Laplacian Estrada index that we had not included in our so-called Group 4). The two large clusters into which this hierarchical clustering can be divided basically divides the complexity measures of the topological descriptors, the network parameters being distributed between both clusters. On the other hand, some topological descriptors were located in the cluster where the complexity measures were found. In this manner, the network parameters Density, Average degree and Average clustering coefficient, and the topological descriptors Normalized edge complexity, Complexity index B and Laplacian Estrada index, were grouped in this cluster. The consistency between these results and the results of the PCA is an indication that the results are solid, reliable and robust.

### 6.4 Network measures

Once the hypothetical tree corresponding to arthropod early evolution has been solved, we will now analyze in detail the results obtained for the different network measures used in this work: network parameters, topological descriptors and complexity measures. For this, we decided to order the groups of arthropods according to the result of the hierarchical clustering, for which the most primitive organisms were located in the center of the figures, to the right the organisms belonging to the right or small evolutionary branch, and to the left the organisms belonging to the left or large evolutionary branch. Thus, the arthropod groups were ordered as follows (from left to right): *Speleonectes*, *Marrella*, *Rehbachiella*, *Triops*, *Canadaspis*, *Yohoia*, *Branchinecta*, *Waptia*, *Olenoides*, *Martinssonia*, *Nebalia* and *Lightiella*.

#### 6.4.1 Network parameters

Ordered according to the early evolutionary bifurcation proposed by PCA and HC, we detected two basic patterns regarding the behavior of the network parameters, which would later be repeated in the other network measures. The first pattern, which we could call *ascending bifurcation*, is characterized by having the minimum value at the hypothesized evolutionary origin and by increasing on both sides of the origin along the right and left evolutionary branches. The second pattern, which we could call *descending bifurcation*, presents the maximum value at the hypothetical evolutionary origin and decreases on both sides of the origin along the right and left evolutionary branches.

The network parameters that behaved according to the first pattern were clearly Diameter, Radius and Average path length, all three members of Group 3 (blue) (Fig. 10). We could also include Nodes and Edges (Group 1, green) within this group, with the difference that in these network parameters the left branch ascends to *Triops* and then descends. On the other hand, the network parameters that behaved according to the second pattern were Density (Group 2, yellow), which had a slight rise in the left branch from *Triops* onwards, and Average degree and Average clustering coefficient (Group 4, orange), despite the fact that these parameters were only higher in *Branchinecta*. As can be seen in the figure, the size of the arthropod networks ranged from approximately 300 nodes (for example, in *Branchinecta*) to 900 nodes (in *Triops*).

**Figure 10.**
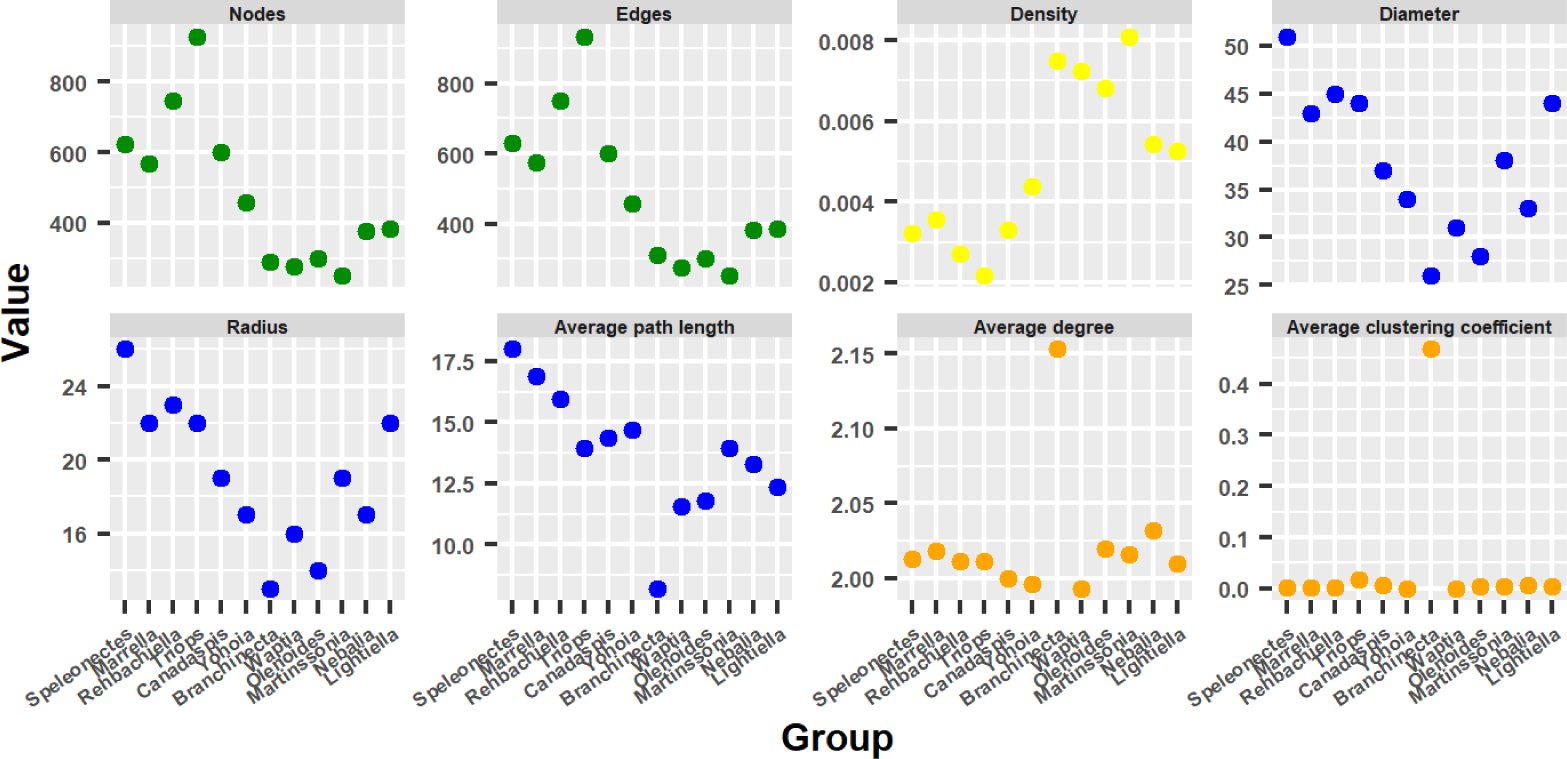
Network parameters. Behavior of network parameters ordering the arthropod networks according to the result of the hierarchical clustering: right branch from the center to the right and left branch from the center to the left. Color represents the group to which each network parameter belongs (see text).

#### 6.4.2 Topological descriptors

Like the network parameters, the topological descriptors showed in their evolution the two basic patterns mentioned above: *ascending bifurcation* and *descending bifurcation* (Fig. 11).

**Figure 11.**
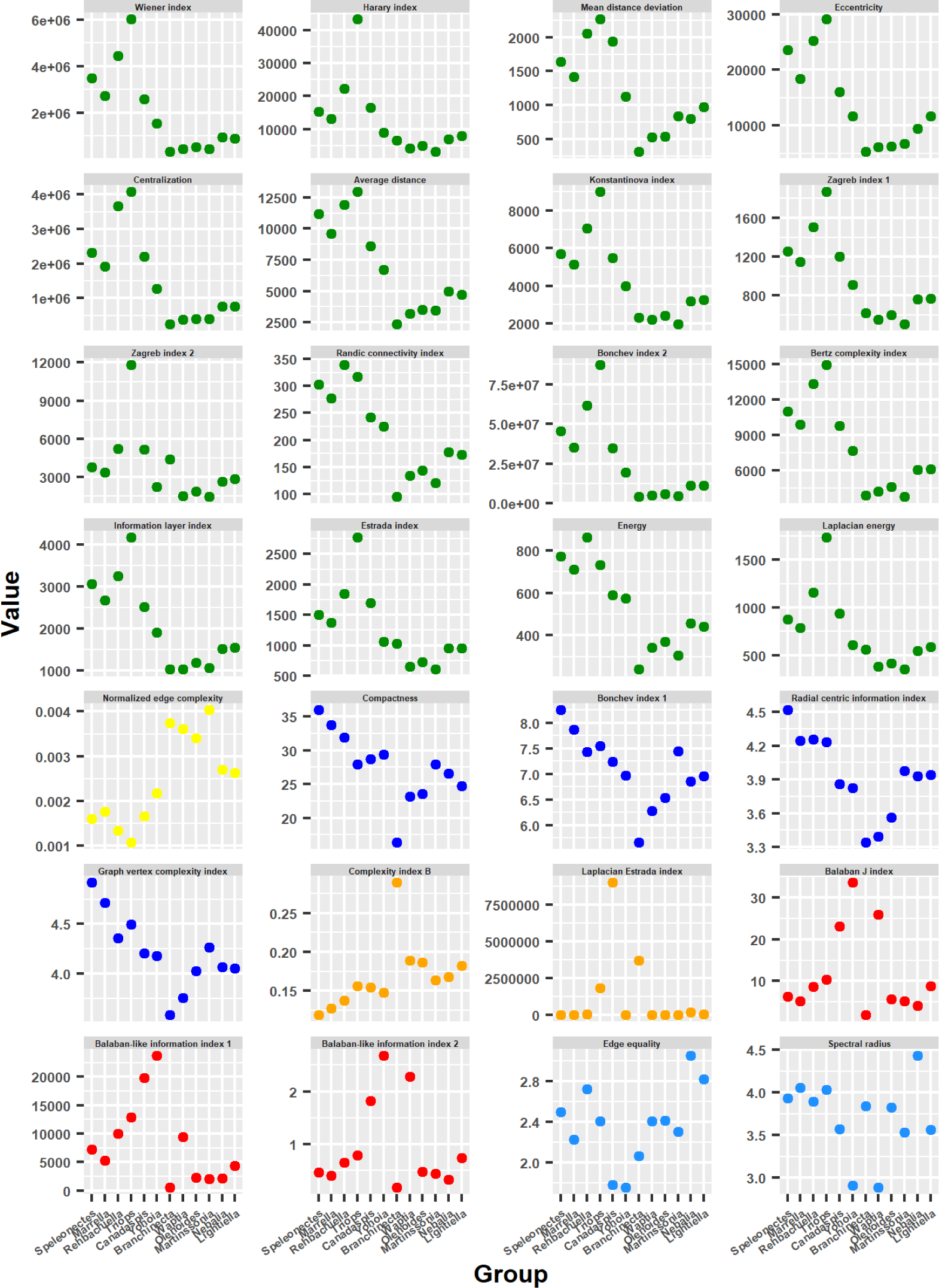
Topological descriptors. Behavior of topological descriptors ordering the arthropod networks according to the result of the hierarchical clustering: right branch from the center to the right and left branch from the center to the left. Color represents the group to which each topological descriptor belongs (see text).

The topological descriptors belonging to Group 1 (green) largely showed the ascending bifurcation pattern: having their minimum values in the central zone, typically in *Branchinecta*, their values ascended in both evolutionary branches. The rise was more pronounced on the left branch, while in general it began to decline from *Triops* onwards, with some exceptions such as Randű connectivity index and Energy. The rise in the right branch was more moderate and variable, with relatively high rises such as in Mean distance deviation, Eccentricity, Randű connectivity index and Energy; and relatively low rises as in Wiener index, Harary index, Centralization and Bonchev index 2. The topological descriptors of Group 3 (blue) behaved similarly to those of Group 1. The most important difference is that in this case there was no descent in the left branch and the ascent in the right branch was more pronounced. Within the topological descriptors that evolved according to the ascending bifurcation pattern, we could also include those of Group 6 (light blue).

On the other hand, the topological descriptors of Groups 2 (yellow), 4 (orange) and 5 (red) evolved following the descending bifurcation pattern. This pattern was especially notable in the topological descriptors of Group 5 (Balaban J index, Balaban-like index 1 and Balaban-like index 2), a group that showed the peculiarity and rarity that one of the groups in which high values were expected of these measures (*Branchinecta*) obtained very low values. Something similar happened with the Laplacian Estrada index (Group 4), in which *Yohoia* obtained much lower values than expected. For its part, Normalized edge complexity (Group 2) behaved almost identically to Density, another member of this group. The maximum value varied in members of these groups. The maximum value in Group 5 was obtained by *Yohoia*. The maximum value in Group 2 was obtained by *Branchinecta*. Meanwhile, the maximum value in Group 4 was obtained by *Branchinecta* (Complexity index B) and *Canadaspis* (Laplacian Estrada index).

#### 6.4.3 Complexity measures

The complexity measures did not behave as clearly as in the previous cases, but even so they could be included in one of the two characteristic evolutionary patterns (Fig. 12).

**Figure 12.**
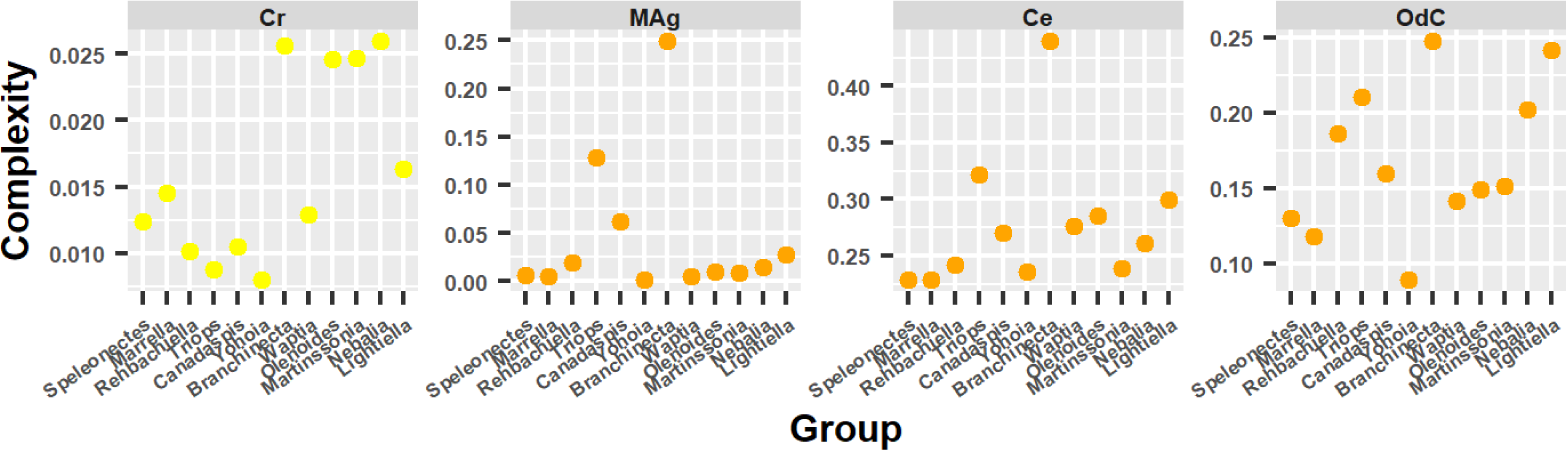
Complexity measures. Behavior of complexity measures ordering the arthropod networks according to the result of the hierarchical clustering: right branch from the center to the right and left branch from the center to the left. Color represents the group to which each complexity measure belongs (see text).

Perhaps the measure that best adapted to one of these two patterns was *MAg* (Group 4), which gave high values for *Canadaspis*, *Branchinecta* and *Triops*, but low for *Yohoia*, a pattern similar to that shown by the Laplacian Estrada index. The same occurred with the complexity measure *Ce* (Group 4), which gave high values only for *Branchinecta*, and intermediate values for *Triops*, a pattern similar to that shown by Complexity index B. In this manner, we could affirm that these two measures had a descending bifurcation pattern, which then seems to be the characteristic evolutionary pattern of the measures of Group 4, as it also seems to be the case for the measures of Group 5. The complexity measure *OdC*, also of Group 4, does not seem to share this characteristic with its group, as it essentially showed an ascending bifurcation pattern, with one exception: *Branchinecta* gave a high value instead of a low one. At the same time, the left branch descended again from *Triops* onwards as it occurred with various topological descriptors that showed this pattern, especially those of Group 1.

On the other hand, the complexity measure *Cr*, belonging to Group 2, had a behavior that was difficult to classify. We could say that it essentially showed an ascending bifurcation evolutionary pattern, with the exception that the measure gave high values for almost all members of the right branch. The values obtained for the case of *Branchinecta* were very high than those expected in the case of an ascending bifurcation pattern. The other two members of this group, Density and Normalized edge complexity, were more easily categorized in the descending bifurcation pattern. The characteristic that *Cr* shared with the other members of Group 2 was that it obtained much higher values for the right branch than for the left branch.

### 6.5 Centrality measures

Now that we have analyzed the behavior of the network measures and the probable early evolutionary process of arthropods, characterized by the presence of a primitive group of arthropods, from which two evolutionary branches arise, right and left, we will begin to try to unravel in what this evolutionary process consists of: what structural changes occur in arthropod networks that explain progress along these evolutionary lines. We will do this by investigating the behavior of centrality measures.

Interestingly, in contrast to what happened with the topological descriptors, most of the centrality measures had a descending bifurcation pattern (red) (Fig. 13). The few exceptions were Betweenness centrality, Information centrality (*netrankr*) and Integration centrality, which had an ascending bifurcation pattern (green) very similar to that obtained by Group 1 of network measures. Within the centrality measures with a descending bifurcation pattern, there were different specific variants. Thus, for example, PageRank centrality and Power centrality (scaled) had a very similar pattern to that obtained by Density and Normalized edge complexity. For its part, Information centrality (*sna*) had a pattern very similar to Complexity index B.

**Figure 13.**
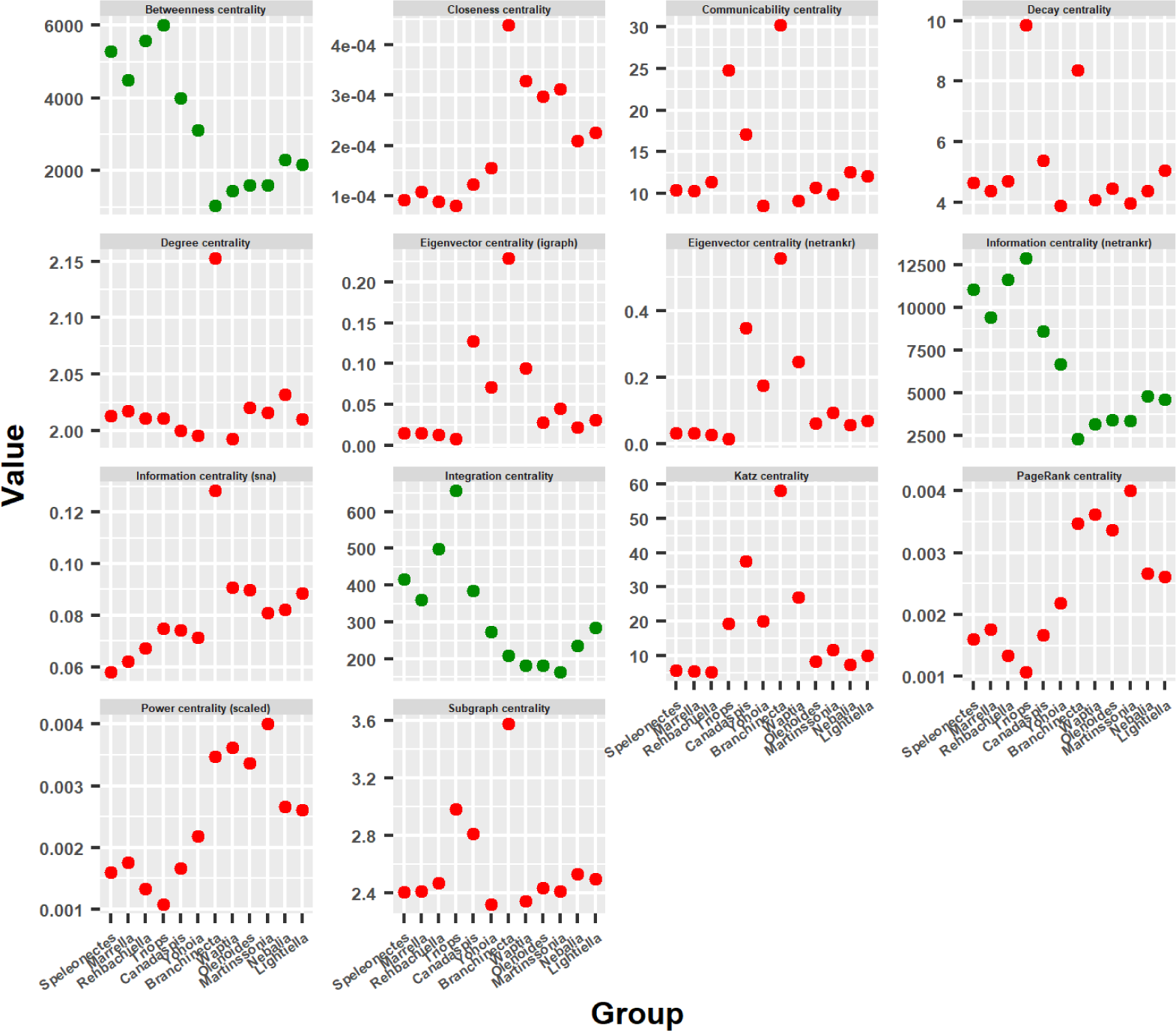
Centrality measures. Behavior of centrality measures’ mean values ordering the arthropod networks according to the result of the hierarchical clustering: right branch from the center to the right and left branch from the center to the left. Most centrality measures had a descending bifurcation pattern (red), while a few exceptions had an ascending bifurcation pattern (green).

The results obtained with the centrality measures are very interesting since they are indicating the presence of a property or characteristic in the most primitive arthropods that is lost throughout the evolutionary process. Among these centrality measures, Eigenvector centrality, Katz centrality, PageRank centrality and Power centrality stand out, since they all derive from the same general basic principle. A general basic principle that will be important for the results and conclusions of our work. We will now study in detail what happens to the centrality measures throughout the evolutionary process.

The most important evidence of the evolutionary process of primitive arthropods seems to be the decoupling between Betweenness centrality (Fig. 14) and Eigenvector centrality (Fig. 15), or what is the same, the disappearance or decline of Eigenvector centrality. While Betweenness centrality remained high and concentrated in the central body axis of all organisms throughout the evolutionary process, both in the left (first and second column) and right (third and fourth column) branches, Eigenvector centrality remained high and concentrated in the central body axis only of the most primitive organisms, specifically the two most primitive organisms of the left and right branch (first and second row), that is, *Yohoia* and *Canadaspis*, and *Branchinecta* and *Waptia*, respectively. In organisms later in the evolutionary process, this centrality measure remained high only in the head or cephalic region. This seems to us the most important result of all the work, since it shows that an important property is lost throughout the evolutionary process, a property that seems to allow or facilitate evolution itself (Eigenvector centrality), at the same time that it is increased or intensified a property that seems rather to slow down and exhaust the evolutionary potential of organisms (Betweenness centrality).

**Figure 14.**
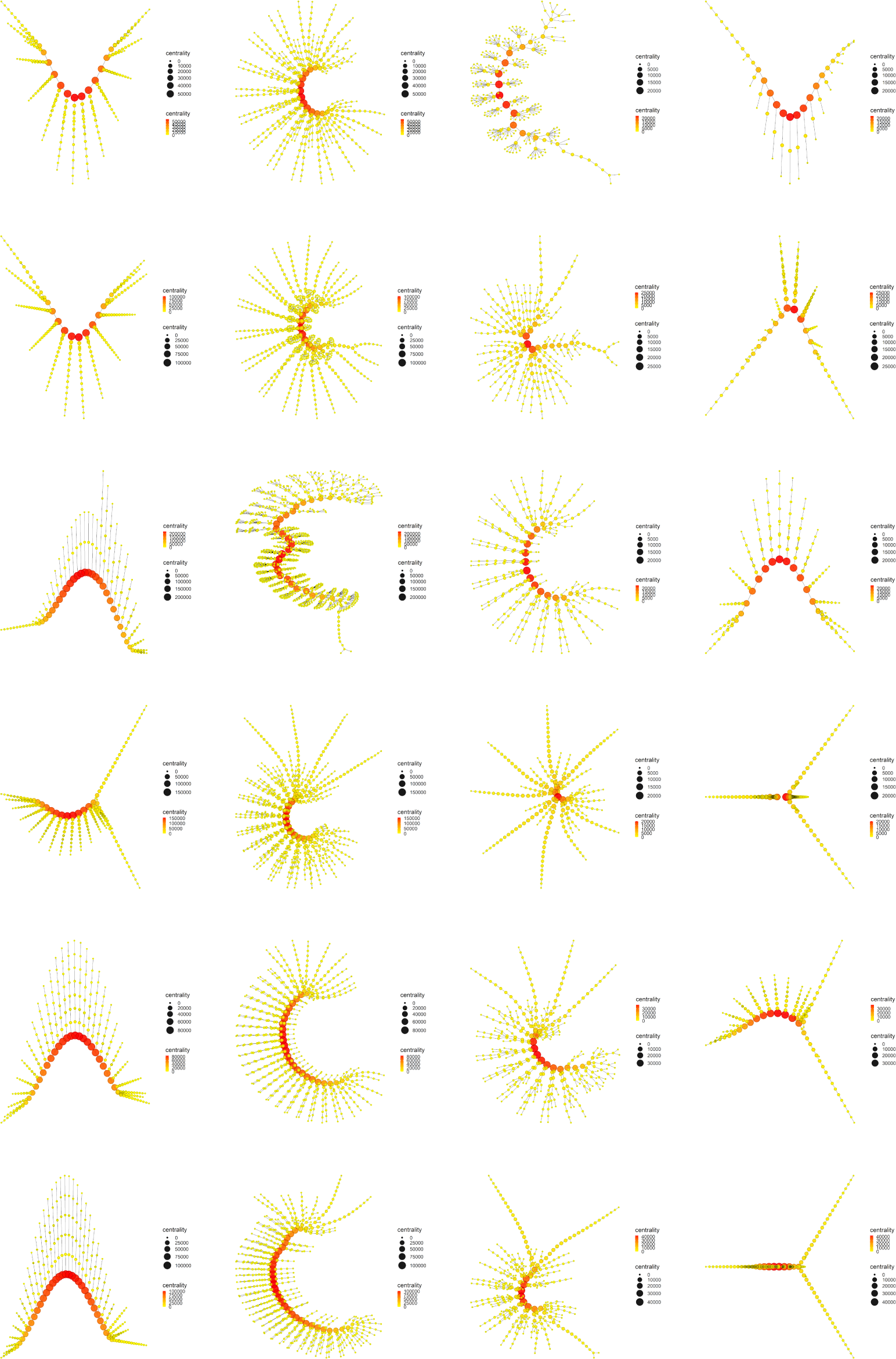
Betweenness centrality. Display and spatialization of arthropod networks’ Betweenness centrality based on two different layout algorithms (Kamada-Kawai (KK) in the two internal columns and MultiDimensional Scaling (MDS) in the two external columns). Arthropod networks are ordered according to the result of the hierarchical clustering. Right branch (column 3 (KK) and 4 (MDS), from row 1 to row 6): *Branchinecta*, *Waptia*, *Olenoides*, *Martinssonia*, *Nebalia*, *Lightiella*. Left branch (column 2 (KK) and 1 (MDS), from row 1 to row 6): *Yohoia*, *Canadaspis*, *Triops*, *Rehbachiella*, *Marrella*, *Speleonectes*. Color (from yellow to red) and size represent the value of the centrality measure of each node (see inset to the right of each network).

**Figure 15.**
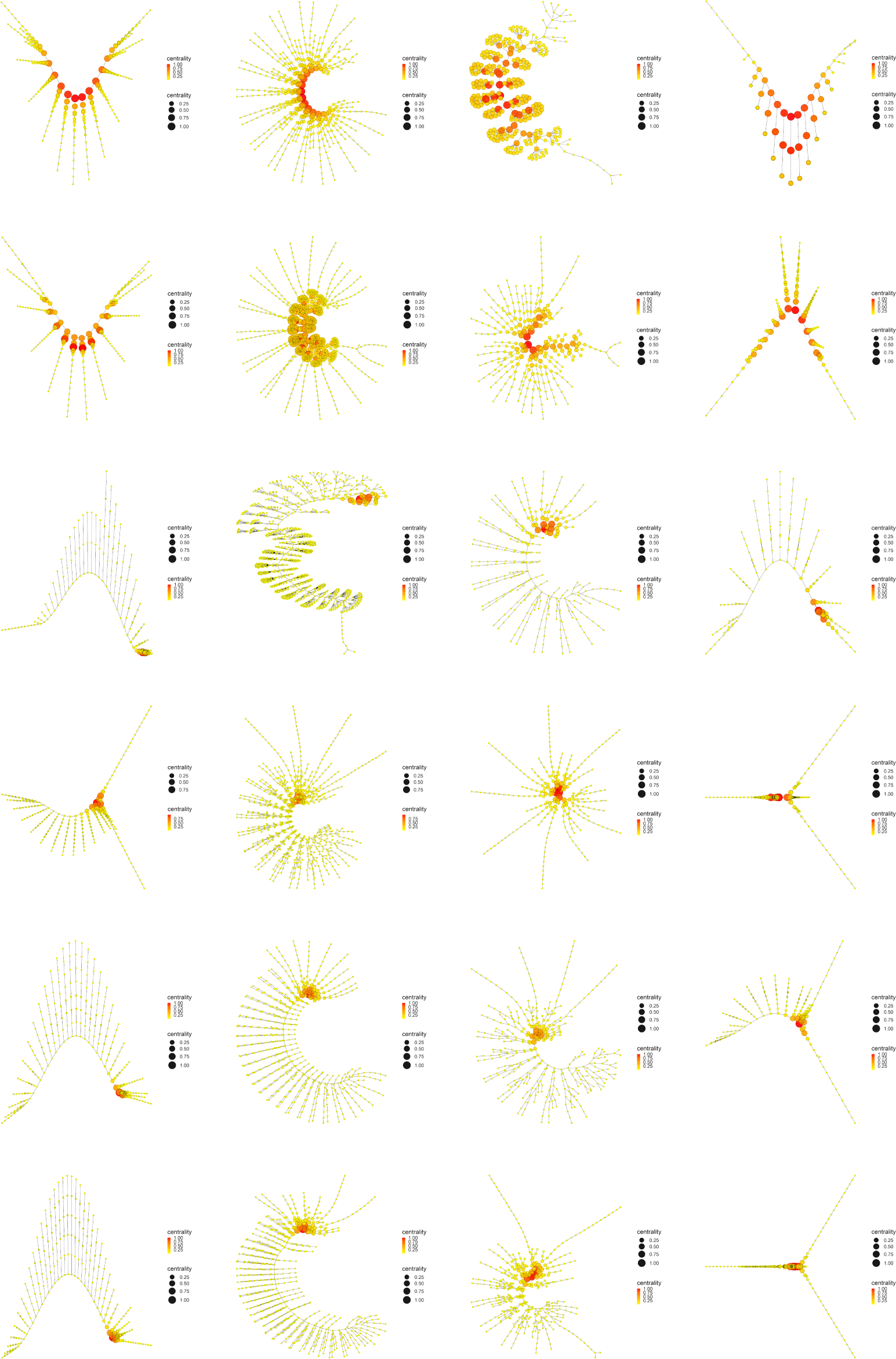
Eigenvector centrality. Display and spatialization of arthropod networks’ Eigenvector centrality based on two different layout algorithms (Kamada-Kawai (KK) in the two internal columns and MultiDimensional Scaling (MDS) in the two external columns). Arthropod networks are ordered according to the result of the hierarchical clustering. Right branch (column 3 (KK) and 4 (MDS), from row 1 to row 6): *Branchinecta*, *Waptia*, *Olenoides*, *Martinssonia*, *Nebalia*, *Lightiella*. Left branch (column 2 (KK) and 1 (MDS), from row 1 to row 6): *Yohoia*, *Canadaspis*, *Triops*, *Rehbachiella*, *Marrella*, *Speleonectes*. Color (from yellow to red) and size represent the value of the centrality measure of each node (see inset to the right of each network).

Something very similar occurred with Katz centrality, a variant of Eigen-vector centrality (Fig. 16). The most important difference in this case was that this centrality measure remained relatively high and concentrated in part of the central body axis of *Triops*, specifically in the first 17 abdominal segments that contain appendages. This can also be verified in Fig. 13. Otherwise, Katz centrality had the same descending behavior throughout the evolutionary process as Eigenvector centrality.

**Figure 16.**
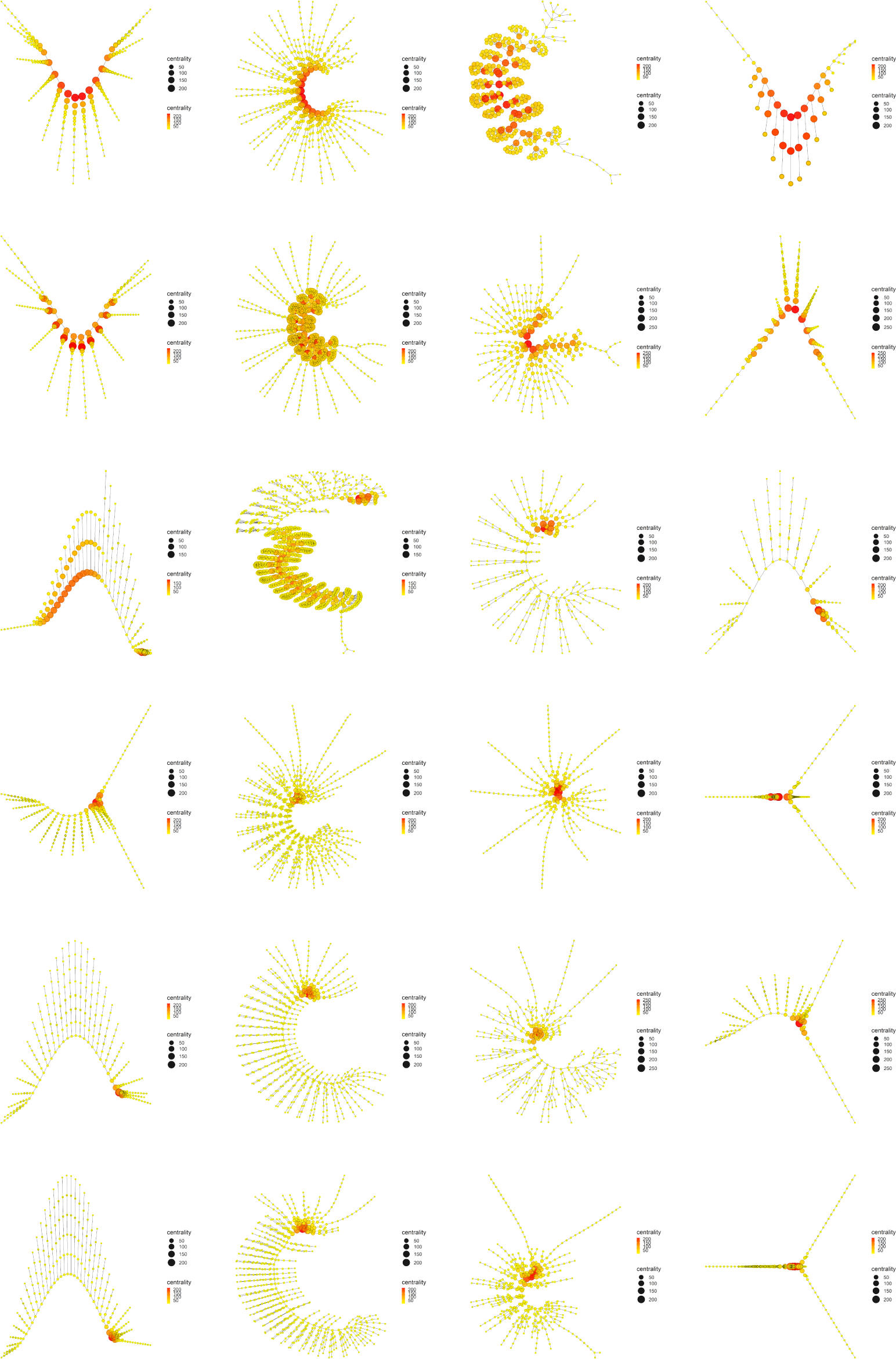
Katz centrality. Display and spatialization of arthropod networks’ Katz centrality based on two different layout algorithms (Kamada-Kawai (KK) in the two internal columns and MultiDimensional Scaling (MDS) in the two external columns). Arthropod networks are ordered according to the result of the hierarchical clustering. Right branch (column 3 (KK) and 4 (MDS), from row 1 to row 6): *Branchinecta*, *Waptia*, *Olenoides*, *Martinssonia*, *Nebalia*, *Lightiella*. Left branch (column 2 (KK) and 1 (MDS), from row 1 to row 6): *Yohoia*, *Canadaspis*, *Triops*, *Rehbachiella*, *Marrella*, *Speleonectes*. Color (from yellow to red) and size represent the value of the centrality measure of each node (see inset to the right of each network).

If we now look at what happened with Power centrality (Fig. 17), what we see is a continuation of the trend followed by Eigenvector centrality and Katz centrality. We see that the centrality measure decreases throughout the evolutionary process, but it persists and lasts longer than the previous measures. In the first three groups (rows) of the left branch (*Yohoia*, *Canadaspis* and *Triops*) and the first two groups (rows) of the right branch (*Branchinecta* and *Waptia*), the values were relatively higher (closer to red) than in the two previous cases. This is particularly noticeable in the case of *Triops*, which obtained intermediate values (orange color) in the body axis in the case of Katz centrality. On the other hand, the Power centrality indices remained at intermediate values in relative terms in the rest of the groups (between dark yellow and light orange), posterior in the two main evolutionary lines. This was particularly noticeable in *Rehbachiella* (fourth group/row of the left branch), and *Martinssonia* and *Lightiella* (fourth and sixth group/row of the right branch), although the effect could also be seen in the rest of the groups.

**Figure 17.**
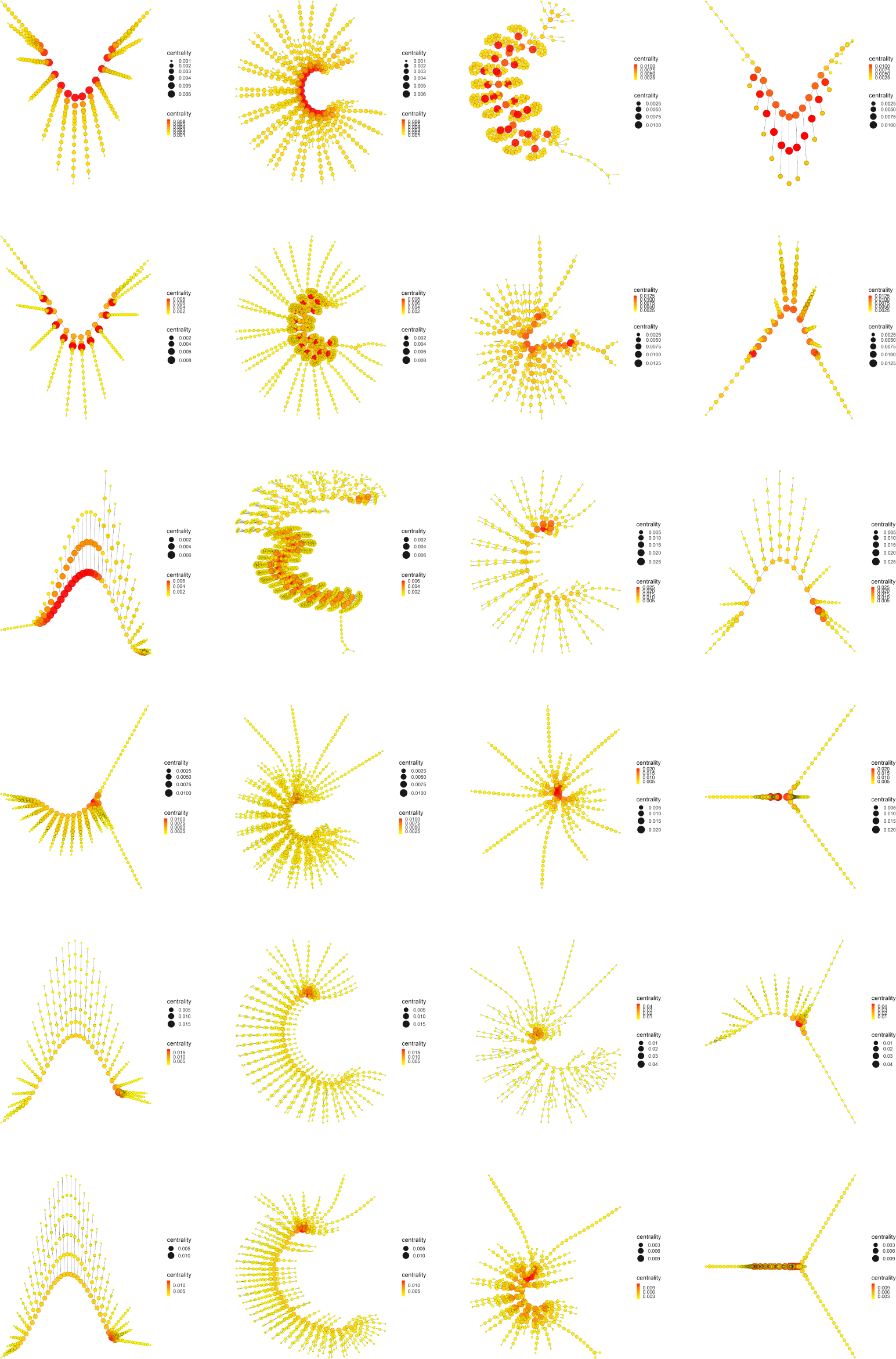
Power centrality. Display and spatialization of arthropod networks’ Power centrality based on two different layout algorithms (Kamada-Kawai (KK) in the two internal columns and MultiDimensional Scaling (MDS) in the two external columns). Arthropod networks are ordered according to the result of the hierarchical clustering. Right branch (column 3 (KK) and 4 (MDS), from row 1 to row 6): *Branchinecta*, *Waptia*, *Olenoides*, *Martinssonia*, *Nebalia*, *Lightiella*. Left branch (column 2 (KK) and 1 (MDS), from row 1 to row 6): *Yohoia*, *Canadaspis*, *Triops*, *Rehbachiella*, *Marrella*, *Speleonectes*. Color (from yellow to red) and size represent the value of the centrality measure of each node (see inset to the right of each network).

This trend was further intensified in the last measure derived from Eigenvector centrality, PageRank centrality (Fig. 18). This measure was highly sensitive to nodes with high degree. Such is the case that in various cases relatively higher values were obtained in these nodes than in the nodes belonging to the central body axis of the organism. The paradigmatic case of this was *Branchinecta*, in which high values were obtained in the basal nodes of the trunk limbs (red color) and intermediate values in the central body axis (light orange color), a circumstance that had not occurred up to now in this organism.

**Figure 18.**
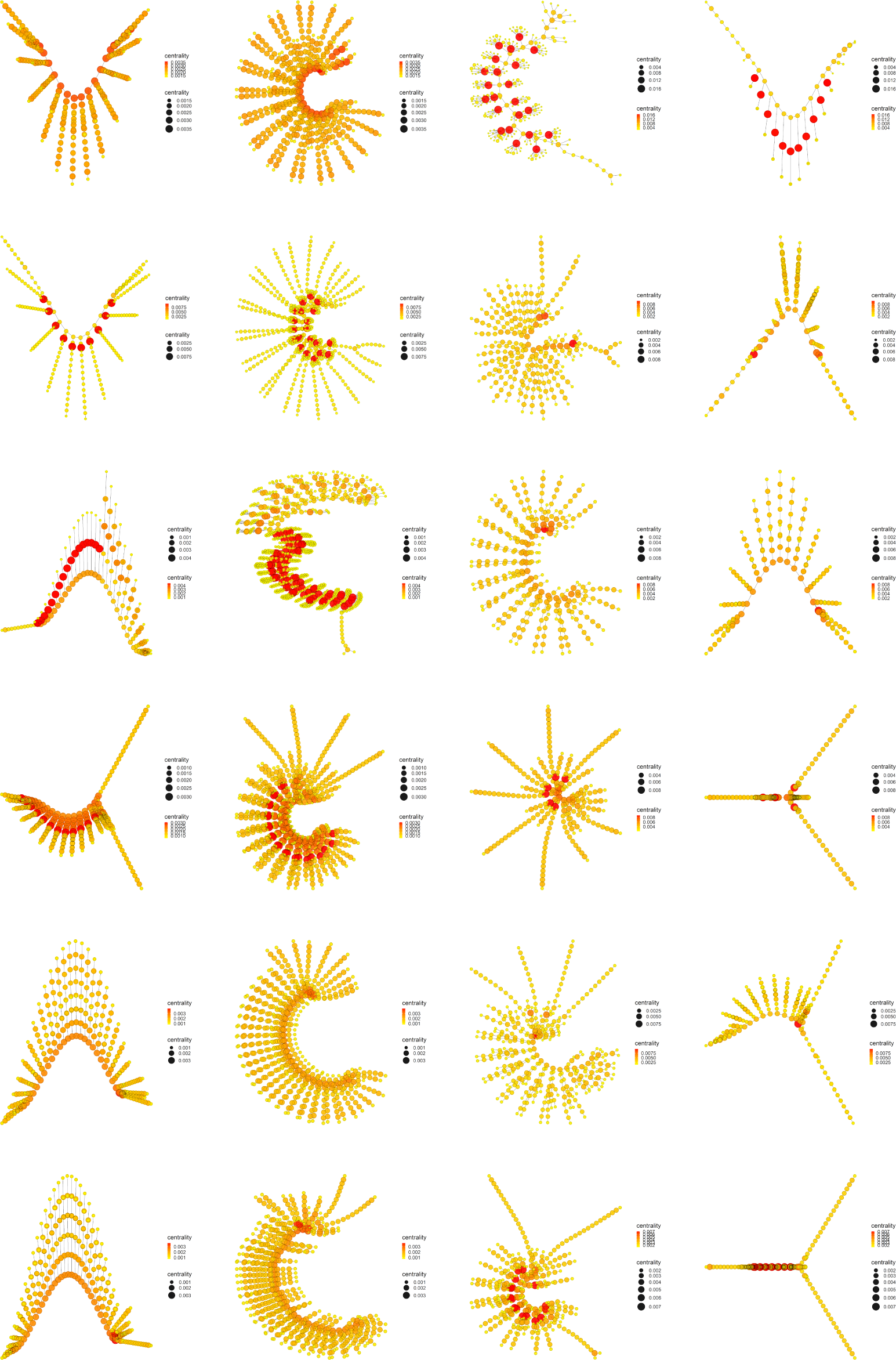
PageRank centrality. Display and spatialization of arthropod networks’ PageRank centrality based on two different layout algorithms (Kamada-Kawai (KK) in the two internal columns and MultiDimensional Scaling (MDS) in the two external columns). Arthropod networks are ordered according to the result of the hierarchical clustering. Right branch (column 3 (KK) and 4 (MDS), from row 1 to row 6): *Branchinecta*, *Waptia*, *Olenoides*, *Martinssonia*, *Nebalia*, *Lightiella*. Left branch (column 2 (KK) and 1 (MDS), from row 1 to row 6): *Yohoia*, *Canadaspis*, *Triops*, *Rehbachiella*, *Marrella*, *Speleonectes*. Color (from yellow to red) and size represent the value of the centrality measure of each node (see inset to the right of each network).

Other interesting results were obtained for Decay centrality (Fig. 19), a measure somewhat related to Closeness centrality. Looking again at Fig. 13, we see that Closeness centrality, along with Power centrality and PageRank centrality, were the measures that showed a remarkably gradual and precise descending bifurcation pattern. Despite this, the Decay centrality pattern did not have these characteristics according to the mean values obtained for these measures. However, visualization of early arthropod networks according to this centrality measure shows that a decrease in the measure also occurs throughout the evolutionary process in both evolutionary branches (Fig. 19). In this sense, it was one of the centrality measures that best showed this decrease in centrality throughout both evolutionary branches, and that at the same time lasted and persisted until the end of them, even more than in Power centrality. Such is the case that relatively high values were obtained even in the last groups of the left branch (*Rehbachiella*, *Marrella* and *Speleonectes*), detecting a decrease from red to dark orange in the nodes corresponding to the central body axis. Something similar occurred in the right branch, in which not only *Branchinecta* and *Waptia* obtained relatively high values in the central body axis (red color), but also *Olenoides* (light red color), descending in *Martinssonia* and *Nebalia* (orange color), and rising again in *Lightiella* (red color), as also verified in Fig. 13.

**Figure 19.**
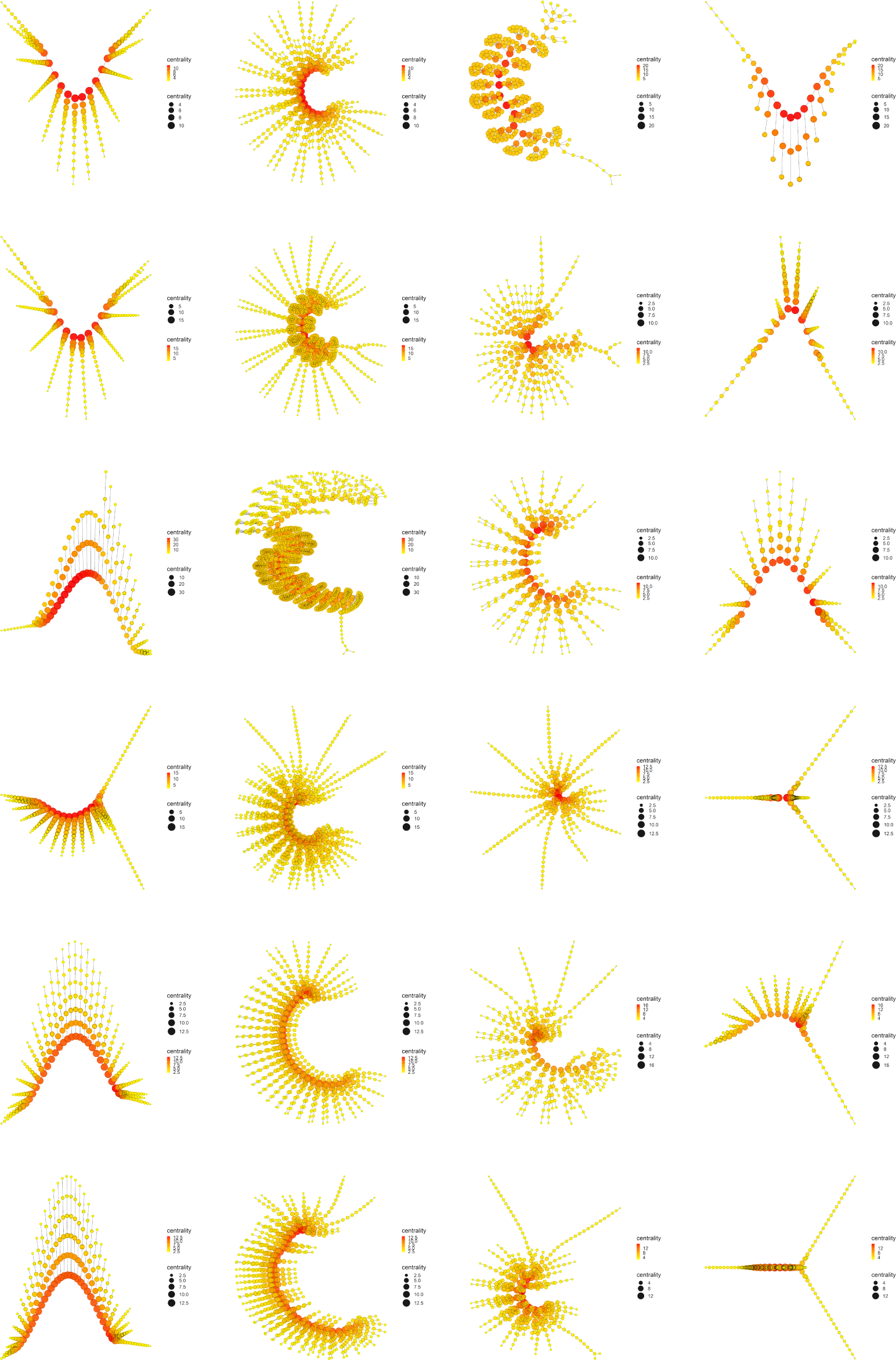
Decay centrality. Display and spatialization of arthropod networks’ Decay centrality based on two different layout algorithms (Kamada-Kawai (KK) in the two internal columns and MultiDimensional Scaling (MDS) in the two external columns). Arthropod networks are ordered according to the result of the hierarchical clustering. Right branch (column 3 (KK) and 4 (MDS), from row 1 to row 6): *Branchinecta*, *Waptia*, *Olenoides*, *Martinssonia*, *Nebalia*, *Lightiella*. Left branch (column 2 (KK) and 1 (MDS), from row 1 to row 6): *Yohoia*, *Canadaspis*, *Triops*, *Rehbachiella*, *Marrella*, *Speleonectes*. Color (from yellow to red) and size represent the value of the centrality measure of each node (see inset to the right of each network).

Finally, the results obtained for Communicability centrality showed a similarity with those obtained for Power centrality (Fig. 20). Perhaps the differences, small and not very noticeable, can be found in the two most primitive organisms of both evolutionary branches, in which Communicability centrality obtained relatively slightly higher values in the nodes corresponding to the central body axis. In everything else, both centrality measures showed highly comparable results, which is interesting for studying the relationship that may exist between them.

**Figure 20:**
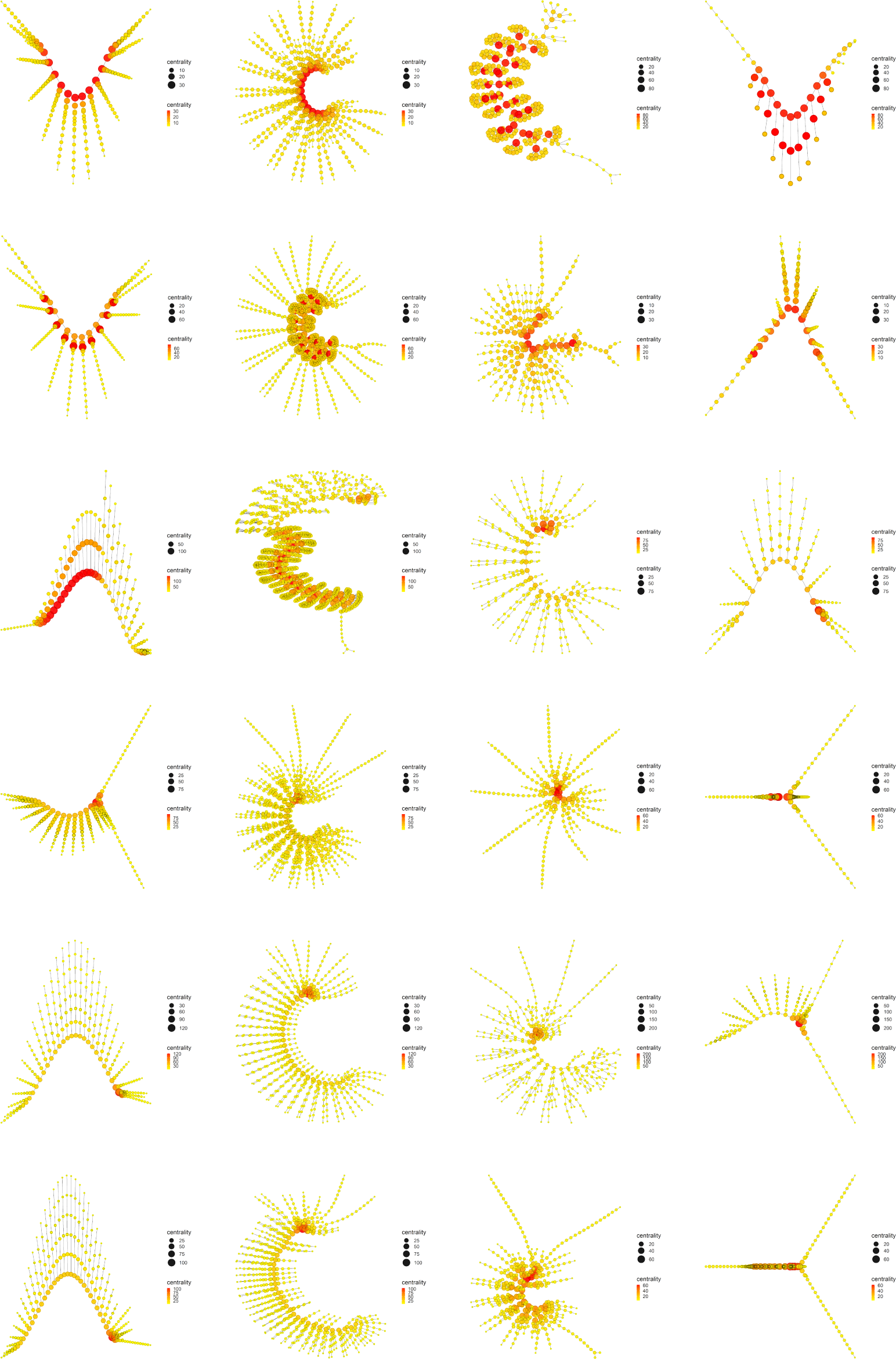
Communicability centrality. Display and spatialization of arthropod networks’ Communicability centrality based on two different layout algorithms (Kamada-Kawai (KK) in the two internal columns and MultiDimensional Scaling (MDS) in the two external columns). Arthropod networks are ordered according to the result of the hierarchical clustering. Right branch (column 3 (KK) and 4 (MDS), from row 1 to row 6): *Branchinecta*, *Waptia*, *Olenoides*, *Martinssonia*, *Nebalia*, *Lightiella*. Left branch (column 2 (KK) and 1 (MDS), from row 1 to row 6): *Yohoia*, *Canadaspis*, *Triops*, *Rehbachiella*, *Marrella*, *Speleonectes*. Color (from yellow to red) and size represent the value of the centrality measure of each node (see inset to the right of each network).

## 7 Discussion

### 7.1 Early arthropod evolution as a process of unfolding of a primitive potential complexity

According to our analysis of the evolutionary process of the early arthropods used in this work, this process is marked by the early bifurcation of two evolutionary branches, which we have called the left or large branch and the right or small branch. In addition to the presence of *Yohoia* and *Canadaspis* as the most primitive members of the left branch, which are somehow organisms generally considered primitive or basal arthropods, the location of *Branchinecta* (Branchiopoda: Anostraca) and *Triops* (Branchiopoda: Notostraca) as the first member of the right branch and the third member of the left branch, respectively, was surprising. This places these organisms as playing a more fundamental and original role than is generally assigned to them. On the other hand, organisms that are generally considered very primitive were located in the last positions of both evolutionary branches. The most paradigmatic case of this was *Marrella*, who ranked as the penultimate representative of the left or large branch.

The evolutionary process as it was ordered and characterized in the present work, using a PCA and Hierarchical clustering, revealed that its main characteristic is given by the presence of a descending bifurcation pattern, in which many of the network measures used in this work decreased as progress was made along both evolutionary branches. The clearest and most important results in this regard were found with the majority of the centrality measures (Fig. 13). Centrality measures such as Eigenvector centrality (Fig. 15), Katz centrality (Fig. 16), Power centrality (Fig. 17) and PageRank centrality (Fig. 18), all based on the same fundamental principle, clearly decreased throughout the evolutionary process. Other centrality measures such as Closeness centrality (Fig. 13), Decay centrality (Fig. 19), related to the previous one, and Communicability centrality (Fig. 20) also decreased. The fundamental difference between them was the speed with which this fall occurred. In this sense, Eigenvector centrality seems to be the earliest detector of primitiveness since only the first two groups of each evolutionary branch had relatively high values of this measure (red color) in the nodes corresponding to the central body axis: *Yohoia* and *Canadaspis* from the left branch, and *Branchinecta* and *Waptia* from the right branch. For their part, Katz centrality, Power centrality and PageRank centrality (in that order) endured and persisted more and more throughout the evolutionary process, the latter being very sensitive to nodes with high degree. In this sense, Decay centrality was the measure that best showed this behavior, being able to detect relatively high values of the measure (dark orange, or even light red) even in the last groups of both evolutionary branches. All this evidences in some way the robustness of the results obtained, especially the structure of the evolutionary tree developed, and reveals an underlying logic in this evolutionary process.

Eigenvector centrality is a centrality measure that measures the influence of a node within the network. Based on the concept that the connection to high-scoring nodes contributes more to the centrality of a node than the connection to low-scoring nodes, this measure assigns relative scores to all nodes in the network. A high score means that one node is connected to many nodes, which in turn are connected to many nodes. In this manner, this centrality measure does not measure the quantity but the quality of the connections. Another way of looking at it is that the Eigenvector centrality is based on the value of the neighbors of a certain entity or node, and not on the intrinsic value of the entity or node itself. This measure is based on the eigenvalue, which means that the value of a node is based on the value of the nodes connected to it: the higher the second, the higher the first. All this leads us to the interpretation that a node with a high Eigenvector centrality is connected to prominent, popular, important nodes, and even if it itself does not have the same importance, it can take advantage of the popularity and influence of its connections. In this sense, a node with a high Eigenvector centrality is a node with many influential ties. Now, how can this interpretation of social networks be translated into the case at hand of a network that represents a morphological structure? What does this “influence” and “popularity” represent in morphological terms? The concept of influence is a concept that can have a translation in terms of morphological evolution and development. Influence somehow represents the power or capacity to control and alter the development of something or someone. In other words, influence represents the power that something or someone has to cause changes in others. A node with high influence, i.e. a node with high Eigenvector centrality, in a network that represents a morphological structure, then represents a node with the power or capacity to cause morphological changes in other nodes, that is, a node that can control and alter the evolutionary development of other nodes. A concept that in this work we are going to call *evolutionary developmental potential*. This concept has a concrete and comparable correlate in developmental biology. There is a concept in this field that is generally called developmental potential, which describes the potential or capacity of a cell or groups of cells to generate and produce different cell types in themselves and in their neighbors. This potential is reduced as the development of the organism progresses. In our case, we apply it to the evolutionary field, although we do not consider it independent and separate from development (hence its name), and we are not applying it to the case of cells or groups of cells but to morphological units or structures.

What conclusions can we then draw from this new concept of evolutionary developmental potential represented and quantified by Eigenvector centrality? The results showed that Eigenvector centrality is higher in the most primitive arthropods and that it is drastically reduced throughout the evolutionary process, both in quantitative terms (normalized mean values, Fig. 13) and in qualitative terms (relative distribution of the centrality measure within each group, Fig. 15). In the most primitive groups (*Yohoia* and *Canadaspis* from the left branch, and *Branchinecta* and *Waptia* from the right branch) the centrality measure was located preferentially in the nodes corresponding to the central body axis, although it was also located in nodes of the limbs or appendages. Let’s move on to investigate this distribution in more detail. In *Yohoia*, the Eigenvector centrality values were relatively high (orange to red) in the nodes corresponding to the body axis segments of the head and trunk, except for the segment corresponding to the eyes (14 segments); and to a lesser extent in the first article of their corresponding appendages, except for the two most distal ones, that is, the great appendage and the trunk appendage 10 (12 pairs of cephalic and trunk appendage basipods). In *Branchinecta*, the Eigenvector centrality values were relatively high (orange to red) in the nodes corresponding to the body axis segments of the trunk (11 segments) and in the nodes corresponding to the base of the trunk limbs (11 limb base pairs). In *Canadaspis*, the Eigenvector centrality values were relatively high (orange to red) in the nodes corresponding to the body axis segments of the trunk and the first and second maxilla (10 segments), and in the first article and the outer ramus lobe from their corresponding appendages (10 pairs of basal articles and 10 pairs of outer ramus lobes). The highest values were found in the outer ramus lobes. Finally, in *Waptia* the Eigenvector centrality values were relatively high (orange to red) in the nodes corresponding to the body axis segments of the cephalothorax and post-cephalothorax (13 segments), and we could also include the first article (podomere) of the post-maxillular appendages 1 to 3 (3 pairs of proximal podomeres). On the other hand, in the rest of the evolutionary series Eigenvector centrality was only detected and concentrated in the cephalic region of the organisms. Thus, for example, in *Triops* the highest value of Eigenvector centrality was found in the carapace (red color), followed by all the nodes to which it was connected: all the cephalic segments (5 segments), including the two eyes (orange color). Something similar occurred in *Rehbachiella*, in which the highest value of Eigenvector centrality was found in the cephalic shield (red color) and, next, the nodes to which it was connected: all the cephalic segments (6 segments, orange color). In general terms, this pattern was repeated in all other organisms: Eigenvector centrality was high first in the cephalic/head shield (red color), and then in the cephalic segments to which it was connected (orange color). In this manner, we can affirm that the evolutionary developmental potential declines throughout the evolutionary process, going from being located along the entire organism body axis, preferably the segments of the head and thorax, to being located only in the cephalic region, cephalic/head shield and connected cephalic segments. This means that primitive organisms have a much more extensive evolutionary-developmental capacity for change and that they have the potential to generate morphological changes along almost their entire body axis (head and thorax): their cephalic and thoracic segments, and the most proximal articles of their appendages, have the capacity to cause morphological changes in themselves and in their neighbors. It is in this sense that these nodes/segments have “influence” in their surroundings. We now turn to Katz centrality, a measure derived from Eigenvector centrality. The main difference between this centrality measure and the previous one is that with this measure *Triops* appears as having high values in its central body axis, something that did not occur with Eigenvector centrality. With this measure, relatively high values of Katz centrality (orange to red color) were observed not only in the cephalic region (carapace, associated cephalic segments and eyes), as was the case with Eigenvector centrality, but also in the body segments corresponding to the abdominal segments with appendages, that is, the first 17 abdominal segments. We could also include the protopods or basal articles of the 17 pairs of abdominal appendages. The rest of the organisms gave a result almost identical to that obtained with Eigenvector centrality. What accounts for this difference between Eigenvector centrality and Katz centrality? Katz centrality is a centrality measure that computes the relative influence of a node by measuring its distance to all other nodes in the network, penalizing each path or connection by an attenuation factor (alpha parameter). Depending on the value of this parameter, this measure can range from Degree centrality (when alpha approaches 0) to Eigenvector centrality (when alpha approaches the inverse of the largest eigenvalue). This can be interpreted as Degree centrality measuring the *local influence* of a node, while Eigenvector centrality measures the *global influence* of a node. In this manner, Katz centrality is a centrality measure that measures both the local and global influence of a node (Zhan, 2017). This is even clearer and more obvious in our case that we calculated Katz centrality without attenuation factor. Following this line of reasoning then, the novelties detected with Katz centrality are due to the fact that this measure is detecting nodes with more local and circumscribed influences than Eigenvector centrality. This means that the evolutionary developmental potential present in the abdominal segments of *Triops* is important, but it has a more limited influence than that of the cephalic segments. Its circle and area of influence to cause morphological changes is smaller and of lesser scope.

If we now study in detail Power centrality, another centrality measure derived from Eigenvector centrality, we see that this trend of higher detection “sensitivity” increases even more. With this measure, the values generally increase in all organisms. In *Triops*, for example, the values obtained for the first 17 abdominal segments are now relatively higher (red color). On the other hand, the already relatively high values in primitive organisms, especially in their central body axes, are further intensified and increased, the most notorious case being the bases of *Branchinecta*’s trunk limbs. However, the most important difference is that organisms that are posterior in the evolutionary process now obtain relatively higher values for this measure. This occurs mainly in *Olenoides*, *Rehbachiella*, *Martinssonia* and *Lightiella*. In *Olenoides*, the segments of the thorax and pygidium (12 segments) now appear in orange. In *Rehbachiella*, the 11 segments of the thorax also appear in orange. In *Martinssonia*, the same occurs with the rest of the cephalic and thoracic segments with appendages (3 segments), and even with the coxa and the base of appendages 2, 3 and 4. For its part, in *Lightiella* the thoracic segments with well-developed appendages (first 7 thoracic segments), and the protopods of the maxillae and thoracopods 1 to 7 (all of 2 articles, except the last of only 1 article), now appear in orange. These results lead us to wonder what Power centrality measures and what “power” means in this context.

Both Eigenvector centrality [38] and Power centrality [39] were developed by the same person. Eigenvector centrality was developed as a centrality measure in which the centrality of a unit consisted of its summed connections to others, weighted by their respective centralities. Power centrality was developed to allow greater flexibility. An extra parameter, attenuation factor or beta parameter, allows to vary the degree of dependency of the score of a unit with respect to the score of the other units. In general terms, this attenuation factor can be interpreted as the probability that a communication or information from one node is transmitted to its neighboring nodes. In this manner, the magnitude of the beta parameter reflects the degree to which the communication is transmitted locally or globally, to the entire structure as a whole. Low values of this parameter predominantly focus and evaluate the local structure, while high values evaluate the position of a node in the structure as a whole. In this sense, the beta parameter can be seen as a radius of influence within which one wants to evaluate the centrality of a node [39]. In this manner, since we used a low value of beta parameter (0.2) we could define that Power centrality is measuring the local evolutionary developmental potential, that is, the power of influence of a morphological unit to cause changes in its immediate surroundings and vicinity. In this case, it makes sense that this centrality measure detects zones of influence not detected by Eigenvector centrality and Katz centrality. We could then define the following scenario: Eigenvector centrality would be revealing long-range zones of influence, Katz centrality would be revealing medium-range zones of influence, and Power centrality would be revealing short-range zones of influence.

These results are highly consistent and point to a scenario in which the evolutionary developmental potential gradually decreases in magnitude and scope throughout the evolutionary process. This means for us the first empirical evidence of our theory of evolution as a process of unfolding [14]. In a previous work, we had found evidence of the presence of such a process in the case of ontogenetic development [15]. In this context, we can affirm that the evolutionary process of early arthropods is largely marked by the decline in the evolutionary developmental potential, mainly measured by Eigenvector centrality, Katz centrality, and Power centrality. This potential unfolds and actualizes in a greater extensive complexity of the organisms, measured by a large part of the topological descriptors (Groups 1 and 3), but also by various network parameters (Groups 1 and 3) and centrality measures (e.g. Betweenness centrality).

An interesting question to ask is whether this evolutionary developmental potential coincides with what we have called intensive complexity in a previous work [15]. It is not a simple question, but we can draw some conclusions based on the obtained results. In the previous work, intensive complexity was represented by the complexity measures, while extensive complexity was represented by the topological descriptors. In this work, the dimension in which these network measures move is mainly given by dimension 1 of the PCA (Fig. 6): topological descriptors are concentrated on the left of the dimensional space (PCA1 negative), and complexity measures are located on the right (PCA1 positive). However, as we have already seen, the temporality of the evolutionary process seems to be given in this analysis predominantly by dimension 3: the most primitive organisms are located in the upper region of the dimensional space (PCA3 positive, Fig. 2). This seems to indicate that the process that we are revealing in this work, which allowed us to define and create the so-called evolutionary developmental potential, introduces a new dimension to the evolutionary developmental process. In this manner, apparently the developmental process and the evolutionary process involve different variables and dimensions in question, although both imply a process of unfolding. This evidently should be further studied in future work.

### 7.2 What is it like to be primitive?

An important question that arises after the results obtained in this work is what it means for an organism to be primitive. More precisely, what morphotopological structure characterizes a primitive organism? Throughout the history of the study of arthropod evolution, the idea that the archetypal primitive arthropod, the *Urarthropod*, was characterized by a simple, monotonous and repetitive structure, has been considered generally accepted. Thus, for example, Hessler & Newman (1975) [9], choosing Cephalocarida (*Hutchinsoniella*), Leptostraca (*Nebalia*) and Notostraca (*Lepidurus*) as representatives of primitive crustaceans, depicted the morphological structure of their urcrustacean as that of an organism consisting of a head and a thorax made up of a repetitive series of about 22 segments, each consisting of a pair of stenopodous limbs of 7 articles. All the appendages of this urcrustacean (mandibles, maxillule 1, maxillule 2 and thoracic limbs) were equal and uniform (Fig. 7 and 8 of the above cited work). One cannot help but see in this reconstruction of the hypothetical urcrustacean, despite all the differences that can be found, the supposedly primitive crustacean that would be discovered a few years later, in 1981, that would be called *Speleonectes* and that would be placed in a new class called Remipedia. At that time, the authors considered that their reconstruction of the hypothetical primitive crustacean had a great similarity and resemblance to trilobites, for which they proposed “A trilobitomorph origin for the Crustacea”. Thus, for these authors, the primitive representative of arthropods in general and crustaceans in particular consisted of an organism with a high degree of serial homology and stenopodous linear limbs. This is the classic and standard view of a primitive arthropod or crustacean. Our work provides clear and compelling evidence against this classical view. All organisms that possessed a morphotopological structure similar to that of this classical view were considered as organisms that were late in the evolutionary process, and organisms in which the evolutionary developmental potential had already decreased substantially, that is, that their potential for change had already been reduced, constrained, and therefore structurally and topologically localized. On the other hand, organisms that were characterized as primitive presented a different morpho-topological structure. They could present a high degree of serial homology, but unlike the classical or standard model, these organisms presented a much more complex segmental topological structure. The segments of these organisms consisted of true *irradiation centers* from the central body segment: a *plexus*. This word is very significant in our context. A plexus is an interwoven network of parts or elements in a structure or system. This expresses very well the type of structure that we are trying to describe. But there are more interesting things in this concept. If one traces the origin of this concept, one finds that a plexus is a structure that is *folded*. We could find here the reason why this structure, the plexus, represents a morphotopological structure of concentrated complexity with the capacity to unfold in a variety of forms thanks to the influential potential that it intrinsically possesses.

There are bibliographic precedents that follow the same line of thought developed and followed in the present work. Olesen, Richter & Scholtz (2001) [40] proposed that segmented trunk limbs have evolved from phyllopodous limbs, based on the study of the embryological development of two branchiopods, one with phyllopodous limbs (*Cyclestheria*) and the other with stenopodous limbs (*Leptodora*). The empirical and logical basis for reaching this conclusion was that both organisms began the development of their limbs in a similar way: “In both species the limbs are formed as ventrally placed, elongate, subdivided limb buds” [40]. This work clearly shows that during its embryological development *Leptodora* goes from having phyllopodous limbs to having stenopodous limbs. The evolutionary translation of this developmental result has the character of inferential, but in any case it is well grounded and justified in the relative phylogenetic positions of both groups. Borradaile (1926) [41] had already speculated about the possibility that the primitive crustacean limb was not stenopodous but phyllopodous, and that articles of stenopodous limbs originated from the endites of phyllopodous limbs. After him, Fryer (1992) [42] proposed a similar hypothesis, pointing out that primitive arthropod appendages did not have stenopodous limbs. The additional contribution of our work regarding this specific point is to provide a theoretical and rational hypothesis about the reason and cause of this evolutionary change: the segmental morpho-topological structure of organisms that have phyllopodous limbs is a structure that contains in itself a higher evolutionary developmental potential that allows it to have the intrinsic capacity to produce morphological changes in its structure and in its more or less neighboring surroundings.

### 7.3 Wonderful Potential Life: early arthropod evolution and the nature of history

In his *Wonderful Life*, Stephen Gould (1989) [43] makes a revision of the Burgess Shale fossils, which date from about 508 million years ago, with the idea of correcting and replacing the iconography of evolution as “cone of increasing diversity”, which he personifies in the figure of Charles Walcott, for the iconography of “decimation and diversification”, which he tries to personify in the figure of Harry Whittington. More profoundly, however, Stephen Gould was not only taking aim against the usual iconography of evolution as a cone or tree of increasing diversity, but also against the view of history as a progressive process of increasing complexity, to propose a model based on *contingency*: “the “pageant” of evolution as a staggeringly improbable series of events, sensible enough in retrospect and subject to rigorous explanation, but utterly unpredictable and quite unrepeatable” [43, p. 14]. Gould posits historical contingency as a third way, a middle way, to the known extremes of historical determinism and complete randomness.

According to Gould, the iconography of the cone made Charles Walcott’s interpretation of Burgess Shale organisms inevitable. These animals were found at the narrow base of the cone, at the origin of pluricellular life, so their diversity must have been limited to a basic morphological simplicity. In this manner, these organisms had to be considered as primitive forms of modern groups, that is, as ancestral forms that, thanks to a temporal increase in complexity, progressed to some modern form. Consequently, Walcott interpreted Burgess Shale organisms as primitive members of known modern arthropod groups. Also according to Gould, the reconstructions of Burgess Shale organisms by Harry Whittington and his colleagues challenged the iconography of the cone and “turned the traditional interpretation on its head” [43, p. 47]. Unlike Walcott, Whittington interpreted Burgess Shale organisms as new morphological groups, not belonging to any modern group. Whittington then would have inverted the cone of life: “The sweep of anatomical variety reached a maximum right after the initial diversification of multicellular animals. The later history of life proceeded by elimination, not expansion. The current earth may hold more species than ever before, but most are iterations upon a few basic anatomical designs” [43, p. 47]. According to this view then, most or all of the *body building plans*, i.e. *Bauplan*, of arthropods appeared at the beginning and not at the end of the evolutionary process: “The maximum range of anatomical possibilities arises with the first rush of diversification” [43, p. 47]. We could rewrite this in this other way: “The greatest *evolutionary developmental potential* is found at the beginning of the evolutionary process”. This was what we found in our work. However, we did not detect an inversion of the cone of life. This may be due to the clustering method used to build the evolutionary tree, which somehow presupposes an arborescent structure. Even so, if we gather all the results obtained in this work, from the PCA to the centrality measures, a clear rationality, directionality and sense of the evolutionary process can be seen, which allows us to think that the results found and organized in the evolutionary tree are very well supported by empirical evidence.

This means that at the beginning of the evolutionary process of arthropods there was the greatest potential for the generation of new body building plans, that is, that primitive arthropods had a greater capacity to generate new forms, and more unique and different forms, than modern arthropods. Here we must also take into account that some of these primitive arthropods with greater potential for change are organisms that still exist today, according to the results of our work. However, the fact that these primitive arthropods have a greater potential does not mean that they are more complex. Rather, they can be quite simple. The problem is what one understands by complex and by simple. This question is rather complex than simple. According to the results of this work, we could say that the simple has a greater potential to generate new forms. However, not just any simple form has a high potential. We have already seen what types of simple structures have a high potential: those that can have serial homology but their segments have a plexus morpho-topology. On the other hand, very complex structures may have a very low potential for change. Their structural elements are so committed, interconnected and integrated into the structure as a whole, that it is very difficult for these structures to change and evolve. This is compatible with Riedl’s concept of *burden* [44].

Returning to Gould, he considers that even the inverted iconography he proposes for the early history of arthropods, based on the Burgess Shale finds, can still be interpreted in the “traditional” terms of evolutionary predictability and directionality: “We can abandon the cone, and accept the inverted iconography, yet still maintain full allegiance to tradition if we adopt the following interpretation: all but a small percentage of Burgess possibilities succumbed, but the losers were chaff, and predictably doomed. Survivors won for cause - and cause includes a crucial edge in anatomical complexity and competitive ability” [43, p. 48]. For Gould, the inverted iconography enables a different alternative that for him was prevented by the iconography of the cone: that survivors did not survive for a justified cause, such as greater morphological complexity, but simply due to mere accidents, such as unpredictable environmental catastrophes. It is in this context that Gould proposes the “experiment”, rather the metaphor, of “replaying life’s tape”: “You press the rewind button and, making sure you thoroughly erase every-thing that actually happened, go back to any time and place in the past - say, to the seas of the Burgess Shale. Then let the tape run again and see if the repetition looks at all like the original. If each replay strongly resembles life’s actual pathway, then we must conclude that what really happened pretty much had to occur. But suppose that the experimental versions all yield sensible results strikingly different from the actual history of life” [43, p. 48]. In the latter case, he would check his hypothesis of historical contingency. This interpretation of Gould has its drawbacks. In the first place, the metaphor of “replaying life’s tape” is confusing, and perhaps the appropriate metaphor would have been that of “time travel”. If life is a tape, then each time we rewind it we will see the reproduction of the same history of life. Now, this metaphor raises various questions: would the contingencies and accidental events that Gould alludes to and resort to be *internal modifications in the content* of the tape or would they rather be *external alterations to the structure* of the tape? In other words, would “impacts of extraterrestrial bodies” produce modifications to the content of the tape or to its structure as a tape itself? It is clear that for Gould the answer is the first option: “any replay of the tape would lead evolution down a pathway radically different from the road actually taken” [43, p. 51]. However, it is clear that Gould adopts an externalist view of life and history in this text, and such a view can only cause external alterations to the tape’s structure and is incapable of generating internal modifications to the tape’s content. The truth is that the iconography, in an increasing or inverted cone, does not change the “view of life” in the sense that he believes: you can have a contingent conception of life with both iconographies. The problem is that in both cases Gould assumes that the shape of the cone modifies and alters the evolutionary process, as if it were a mold to which the evolutionary process must conform. Our work brings a new vision to life evolution, which is even more faithful to the idea of “life’s tape”. Our proposal is that life has a potential that unfolds and actualizes in what we call history. The potentiality of life would be the tape’s content, and its reproduction would be history itself. The results of our work provide evidence that life evolution has sense and directionality, that is, that it follows a path marked by intrinsic internal patterns, which can be studied and revealed, and could even be predicted.

## 8 Acknowledgments

## 8.2 Author contributions

A.O. carried out all the steps that led to the writing of this manuscript: he conceived the project, acquired and analyzed the data, obtained the results, made the figures, reached the conclusions, wrote and revised the resulting manuscript.

## 8.4 Data availability

All data needed to evaluate the conclusions in the paper are present in the paper and/or the Supplementary Materials.

